# Multi-trait genome-wide association meta-analysis of dietary intake identifies new loci and genetic and functional links with metabolic traits

**DOI:** 10.1101/623728

**Authors:** Jordi Merino, Hassan S. Dashti, Chloé Sarnowski, Jacqueline M. Lane, Miriam S. Udler, Petar V. Todorov, Yanwei Song, Heming Wang, Jaegil Kim, Chandler Tucker, John Campbell, Toshiko Tanaka, Audrey Y. Chu, Linus Tsai, Tune H. Pers, Daniel I. Chasman, Josée Dupuis, Martin K. Rutter, Jose C. Florez, Richa Saxena

**Affiliations:** Center for Genomic Medicine, Massachusetts General Hospital, Boston, MA, USA; Programs in Metabolism and Medical & Population Genetics, Broad Institute of MIT and Harvard, Cambridge, MA, USA; Diabetes Unit, Massachusetts General Hospital, Boston, MA, USA; Department of Medicine, Harvard Medical School, Boston, MA, USA; Department of Anesthesia, Critical Care and Pain Medicine, Massachusetts General Hospital and Harvard Medical School, Boston, MA, USA; Department of Biostatistics, Boston University School of Public Health, Boston, MA, USA; Novo Nordisk Foundation Centre for Basic Metabolic Research, Faculty of Health and Medical Sciences, University of Copenhagen, Copenhagen, Denmark; Division of Sleep and Circadian Disorders, Department of Medicine, Brigham and Women’s Hospital, Boston, MA, USA; Division of Sleep Medicine, Harvard Medical School, Boston, MA, USA; Division of Endocrinology, Diabetes and Metabolism, Department of Medicine, Beth Israel Deaconess Medical Center, Harvard Medical School, Boston, MA, USA; Department of Biology, University of Virginia, Charlottesville, VA, USA; Translational Gerontology Branch, National Institute on Aging, Baltimore, MD, USA; Merck & Co., Inc., Boston, MA, USA; Department of Epidemiology Research, Statens Serum Institut, Copenhagen, Denmark; Division of Preventive Medicine, Brigham and Women’s Hospital and Harvard Medical School, Boston, MA, USA; Division of Genetics, Brigham and Women’s Hospital and Harvard Medical School, Boston MA, USA; Division of Endocrinology, Diabetes & Gastroenterology, School of Medical Sciences, Faculty of Biology, Medicine and Health, University of Manchester, UK; Manchester Diabetes Centre, Manchester University NHS Foundation Trust, Manchester Academic Health Science Centre, Manchester, UK

## Abstract

Dietary intake, a major contributor to the global obesity epidemic^1–5^, is a complex phenotype partially affected by innate physiological processes.^6–11^ However, previous genome-wide association studies (GWAS) have only implicated a few loci in variability of dietary composition.^12–14^ Here, we present a multi-trait genome-wide association meta-analysis of inter-individual variation in dietary intake in 283,119 European-ancestry participants from UK Biobank and CHARGE consortium, and identify 96 genome-wide significant loci. Dietary intake signals map to different brain tissues and are enriched for genes expressed in β1-tanycytes and serotonergic and GABAergic neurons. We also find enrichment of biological pathways related to neurogenesis. Integration of cell-line and brain-specific epigenomic annotations identify 15 additional loci. Clustering of genome-wide significant variants yields three main genetic clusters with distinct associations with obesity and type 2 diabetes (T2D). Overall, these results enhance biological understanding of dietary composition, highlight neural mechanisms, and support functional follow-up experiments.

---

As dietary components are strongly correlated, we conducted a multi-trait genome-wide association meta-analysis of overall variation in dietary intake among 283,119 European-ancestry participants from the UK Biobank^15^ and the Cohorts for Heart and Aging Research in Genomic Epidemiology (CHARGE) Consortium^14^ (Methods; Supplementary Table 1). First, we conducted single-trait GWAS for the proportion of total energy intake from carbohydrate, fat, and protein in UK Biobank (n=192,005). Next, single-trait GWAS from the UK Biobank and CHARGE Consortium (n=91,114) were meta-analyzed and combined into a multi-trait genome-wide association meta-analysis (Methods). An analysis overview is presented in Supplementary Fig. 1.

We evaluated dietary intake using 24-hour web-based diet recall in the UK Biobank^16,17^ and validated food frequency questionnaires, diet history and diet records in the CHARGE Consortium.^14^ We observed strong genome-wide genetic correlations for nutrient estimates between the UK Biobank and CHARGE datasets (*r*_g_ >0.6 for all; *P* <0.001; Supplementary Table 2). The quantile-quantile plots of single-trait and multi-trait meta-analyses showed moderate inflation (λ_GC_ ranging from 1.12 to 1.17) with a linkage disequilibrium (LD) score intercept^18^ of ~1 (standard error (s.e.) = 0.01), indicating that most inflation could be explained by polygenic signal (Supplementary Fig. 2, Supplementary Table 3). In single-trait meta-analyses, genome-wide SNP-based heritability^19^ was estimated at 3.9% (s.e.=0.01), 2.8% (s.e.=0.01), and 3.0% (s.e.=0.01) for carbohydrate, fat, and protein, respectively (Supplementary Table 3), in line with previous GWAS findings^12,14^ and other behavioral phenotypes such as tobacco or alcohol use.^20^

In the multi-trait meta-analysis, 156 lead variants in 96 distinct genomic loci reached genome-wide significance (Methods, Fig. 1a, Supplementary Dataset, Supplementary Tables 4,5). Single-trait meta-analyses of carbohydrate, fat, and protein intake identified 10, 8, and 9 genome-wide significant loci, respectively (Supplementary Tables 6-8). To account for potential reverse causation effects, we investigated whether individuals at extreme quartiles of genetic risk for body mass index (BMI), T2D, or coronary artery disease (CAD) reported different dietary composition (Methods). We found no evidence of such effects, except for individuals with higher number of BMI risk alleles that reported higher protein intake (Supplementary Table 9). In BMI-adjusted multi-trait meta-analysis, 88 out of 96 identified signals retained genome-wide significance (Supplementary Table 10).

**Fig. 1.**
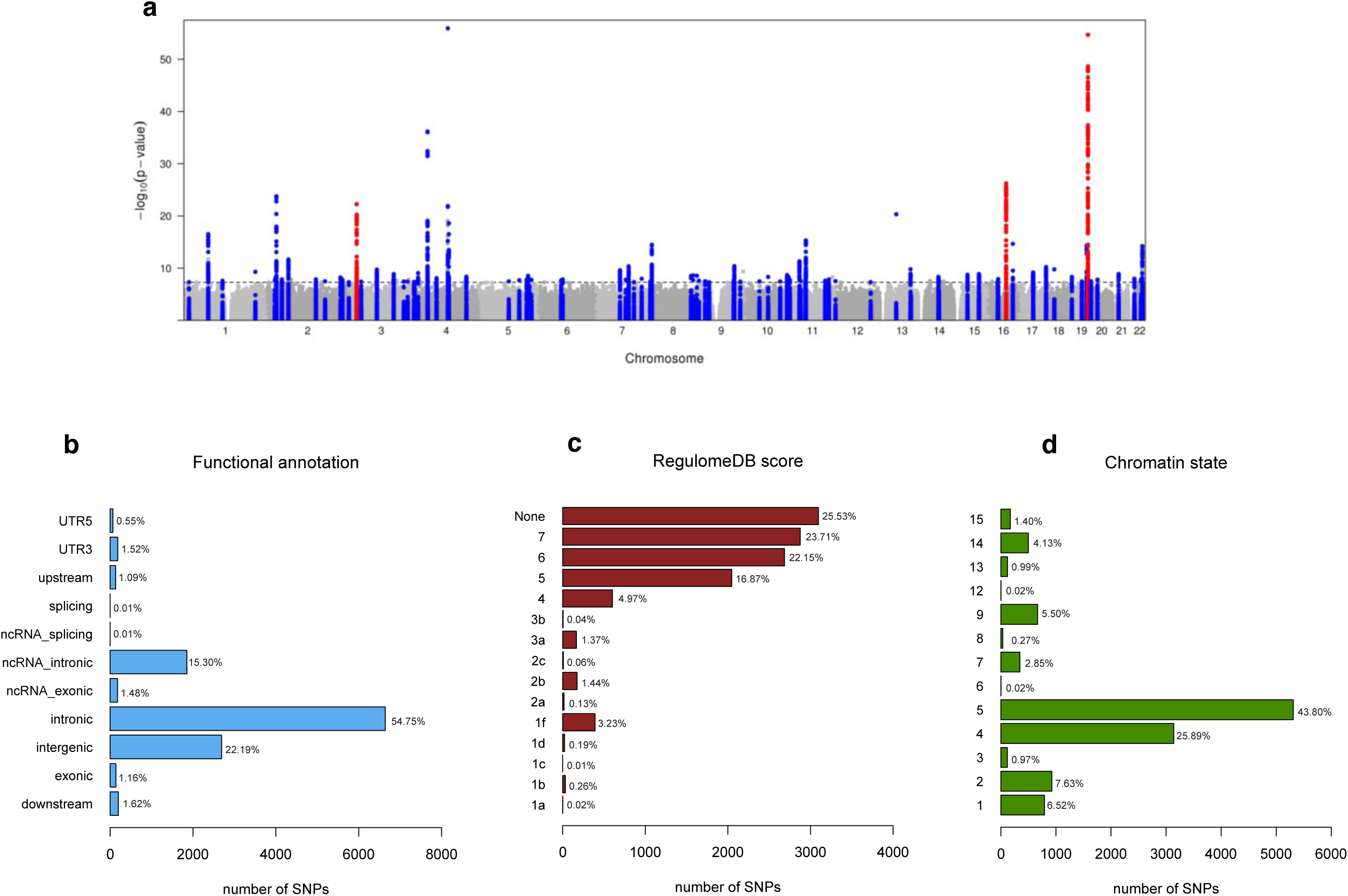
SNP-based association with dietary composition in the multi-trait genome-wide association meta-analysis of 283,119 individuals. **a)** Manhattan plot shows the −log_10_ *P* values (y-axis) for all genotyped and imputed SNPs passing quality control in each GWAS, plotted by chromosome (x-axis). Horizontal line denotes genome-wide significance (*P* =5 × 10^−8^). Red denotes previously identified variants, and blue novel variants. **b)** Distribution of the functional consequences of SNPs in genomic risk loci in the meta-analysis. **c)** Distribution of the RegulomeDB scores for each SNP in genomic risk loci, with a low score indication a higher likelihood of the SNP having a regulatory function (Methods). **d)** The minimum chromatin state across 127 tissues and cell types for SNPs in genomic risk loci, with states 1-7 referring to open chromatin states (Methods).

Functional annotation of all candidate variants in associated loci (*n*=12,675 variants; Methods) identified 67 nonsynonymous variants among them (Fig. 1, Supplementary Tables 5, 11), five of which were the lead variants at their respective locus (Table 1). Several genes in Table 1 have been previously associated with dietary phenotypes, neurological processes, and cardiometabolic risk factors (Supplementary Table 12). For example, the nonsynonymous variant rs4975017 in *KLB*, which encodes the β-Klotho, a primary ‘zip code’-like receptor for endocrine fibroblast growth factor 21 (FGF21) signaling, may impair the affinity between β-Klotho and FGF21^21^. FGF21’s relevance for eating behavior has enabled the development of a drug found to reduce sugar intake and sweet taste preference when administered to monkeys and humans with obesity.^22,23^ A common missense mutation in rs601338 results in a premature stop codon in the exon 2 of the fucosyltransferase 2 (*FUT2*) gene. *FUT2* regulates expression of H group antigens on the surface of epithelial cells,^24^ but also affects vitamin B12 and folate pathways,^25^ and is associated with multiple complex phenotypes such as inflammation,^26,27^ cognitive decline,^28^ and lipids levels.^29^

**Table 1.** Non-synonymous lead variants associated with dietary intake in a multi-trait meta-analysis of dietary intake in 283,119 individuals.

| Locus | CHR | BP | SNP | Gene | AA change | EA | NEA | EAF | P<sub>multi-trait</sub> | CHO<br>β (SE) % | FAT<br>β (SE) % | PROT<br>β (SE) % |
| --- | --- | --- | --- | --- | --- | --- | --- | --- | --- | --- | --- | --- |
| 7 | 2 | 27851918 | rs3749147 | *GPN1* | Arg12Lys | A | G | 0.25 | 5.55E-11 | 0.02(0.02) | 0.07 (0.02) | 0.04 (0.01) |
| 24 | 4 | 39450229 | rs4975017 | *KLB* | Gln1020Lys | A | C | 0.33 | 8.61E-11 | 0.07 (0.02) | 0.01 (0.02) | 0.03 (0.01) |
| 27 | 4 | 100239319 | rs1229984 | *ADH1B* | His8Pro | T | C | 0.04 | 1.23E-56 | 0.11 (0.07) | 0.54 (0.05)** | 0.19 (0.03)** |
| 28 | 4 | 103188709 | rs13107325<sup>a</sup> | *SLC39A8* | Ala391Thr | T | C | 0.08 | 2.54E-19 | 0.13 (0.04) | -0.07 (0.03) | 0.10 (0.02)** |
| 38 | 7 | 73012042 | rs35332062<sup>a</sup> | *MLXIPL* | Ala358Val | A | G | 0.13 | 6.03E-09 | 0.12 (0.03) | -0.10 (0.03) | -0.06 (0.01)* |
| 64 | 11 | 47701528 | rs12286721<sup>a</sup> | *AGBL2* | Met615Ile | A | C | 0.55 | 7.28E-09 | 0.07 (0.02) | -0.01 (0.01) | 0.03 (0.01) |
| 73 | 14 | 60903757 | rs1254319 | *C14orf39* | Leu524Phe | A | G | 0.30 | 4.77E-09 | 0.09 (0.02) | -0.09 (0.02)* | 0.02 (0.01) |
| 82 | 17 | 44060775 | rs63750417<sup>a</sup> | *MAPT* | Pro202Leu | T | C | 0.22 | 1.29E-08 | 0.17 (0.03)* | 0.07 (0.01) | -0.04 (0.01) |
| 87 | 19 | 45411941 | rs429358 | *APOE* | Cys130Arg | T | C | 0.85 | 4.48E-15 | -0.20 (0.03)** | 0.15 (0.03)** | 0.02 (0.01) |
| 88 | 19 | 49206674 | rs601338<sup>a</sup> | *FUT2* | Trp154Ter | A | G | 0.49 | 2.12E-30 | 0.15 (0.02)** | -0.16 (0.02)** | -0.06 (0.01)** |
| 96 | 22 | 41604353 | rs20551<sup>a</sup> | *EP300* | Ile971Val | A | G | 0.72 | 1.91E-8 | -0.09 (0.02)* | 0.06 (0.02) | -0.02 (0.01) |

Note: The lead variants in approximately 5% of loci were non-synonymous. Shown here are all non-synonymous variants, and selected non-synonymous variants in high LD with a lead variant for a given genomic loci (The complete list of 67 non-synonymous variants (38 reaching genome-wide significance) is detailed in Supplementary Table 6). <sup>a</sup> These variants were not themselves lead variants, but were in high LD with a sentinel variant ( $r^2 > 0.8$ from the 1000 Genomes European population). P<sub>multi-trait</sub> value derived from the multi-trait genome-wide association meta-analysis. Effect sizes (β and SE) derived from single-trait GWAS meta-analysis for each respective macronutrient. CHO, carbohydrate; PROT, protein. \* $P < 5 \times 10^{-5}$ , \*\* $P < 5.00 \times 10^{-8}$ in the single-trait genome-wide association meta-analysis. CHR, chromosome; BP, Base Pair based on Hg 19; EA, Effect Allele; NEA, Non-Effect Allele; EAF, Effect Allele Frequency.

We next tested dietary intake signals for enrichment of gene expression in 53 tissue types obtained from the Genotype-Tissue Expression (GTEx) Project^30^ and in 10,651 predefined gene sets derived from MSigDB^31^ (Methods). Signals were significantly enriched for genes predominantly expressed in several brain tissues including the cerebellum (*P*=1.1×10^−6^), frontal cortex (*P*=2.9×10^−5^), anterior cingulate cortex (*P*=1.3×10^−4^), nucleus accumbens (*P*=2.6×10^−4^), and hypothalamus (*P*=9.5×10^−4^) (Fig. 2, Supplementary Table 13). These findings support the notion that dietary intake is, at least, partially controlled by the central nervous system including areas that are known to influence energy homeostasis and appetite control. Two gene sets were significantly enriched for association with dietary intake including neurogenesis (*P*=2.6×10^−7^), and nuclear translocation (*P*=2.3×10^−6^) (Fig. 2, Supplementary Tables 14-15).

**Fig 2.**
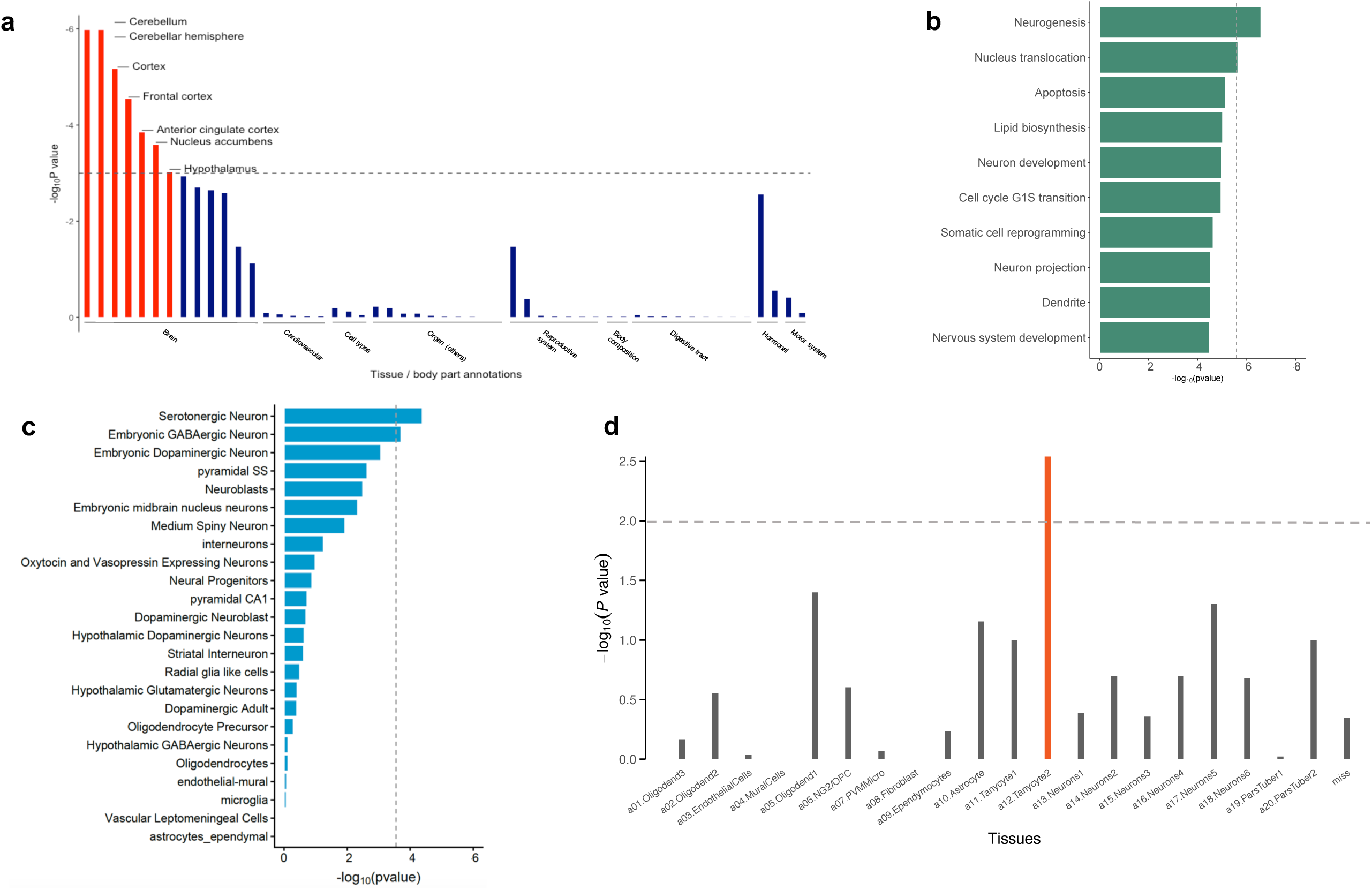
Implicated tissue, pathways, and cell expression profiles for dietary intake in the multi-trait genome-wide association meta-analysis of 283,119 individuals. **a)** MAGMA tissue expression enrichment analysis using gene expression per tissue based on GTEx RNA-seq data for 53 specific tissue types for multiple-trait macronutrient intake. Expression values (Reads Per Kilobase Million – RPKM) were log2 transformed with winsorization at 50 and averaged per tissue (Methods). The dashed line indicates the threshold for significance (P<0.05/53). **b)** Gene set enrichment analysis in 10,651 curated gene sets representing known biological functions and metabolic pathways derived from MSigDB. Only shown the top 10 findings. The dashed line indicates the threshold for significance (P<0.05/10,651 gene sets). **c)** Single-cell expression analysis of genes related to dietary intake in 24 cell types from mouse brain (Methods). The x axis shows the –log_10_ transformed two-tailed P value of association of multi-trait genome-wide association meta-analysis statistics with cell-specific gene-expression levels in a linear model. The dashed grey line indicates the Bonferroni-corrected significance threshold (P<0.05/24 level 1 + 149 level 2 annotations). **d)** Single-cell expression analysis of genes related to dietary intake using Drop-seq data from 20,921 individual cells in and around the adult mouse Arc-ME (Methods). The x axis shows the –log_10_ transformed two-tailed P value of association of multi-trait genome-wide association meta-analysis statistics with cell-specific gene-expression levels in a linear model. The dashed grey line denotes FDR=0.05 derived by DEPICT.

To evaluate whether the genomic loci implicated in dietary intake map onto specific brain cell types, we used two independent brain single-cell RNA sequencing datasets (Methods).^32,33^ Cell-type-specific gene expression in 24 mouse-derived brain cell types showed a significant enrichment for dietary intake genomic loci in genes expressed in serotonergic (*P*=4.1×10^−5^) and embryonic GABAergic neurons (*P*=1.9×10^−4^) (Fig. 2, Supplementary Table 16). Given that several subtypes of serotonergic and embryonic GABAergic neurons have been recently identified in and around the adult mouse hypothalamic arcuate-median eminence complex nucleus (Arc-ME),^10,32^ we used cell-type-specific gene expression in 50 transcriptionally distinct cell populations in the Arc-ME (Methods). We identified enrichment of dietary-linked gene expression in β1 tanycytes (*P*=2.9×10^−3^), but not in other neuronal subpopulations (Fig. 2d, Supplementary Table 17). Tanycytes are a specialized type of ependymal cell lining the wall of the third ventricle with a variety of functions including neurogenesis or controlling the chemical exchange between brain parenchyma, cerebrospinal fluid, and bloodstream.^34–36^ In addition to the dietary intake associated signals, β1 tanycytes were enriched for the expression of genes previously associated with lipids (*ABCG5, APOB*) and growth factor (*FGF18*) (Supplementary Table 17).

We next investigated whether the integration of genomic annotations from several cell lines and brain tissues could identify additional relevant genetic associations influencing dietary intake. We used fGWAS,^37^ a hierarchical modeling approach to re-weight association estimates by using information from the most relevant functional annotations (Methods). By combining separate statistically significant annotations and applying model selection and cross-validation, we showed that the most relevant annotations for dietary intake were DNase I hypersensitivity sites in fetal brain (log_2_ enrichment of 1.22, 95%CI: 0.28 to 1.93), repressed chromatin in B cells (−0.71, 95%CI: −1.22 to −0.17), and weak enhancers in epithelial cells (2.50, 95%CI: 1.23 to 3.39) (Supplementary Tables 18-19). We then used parameters from this model as priors to re-weight the multi-trait meta-analysis, and identified 15 additional genomic regions with high-confidence associations (posterior probability of association [PPA] > 0.8) only when the model incorporated information from functional annotations (Fig. 3, Supplementary Table 20). These additional signals comprised sub genome-wide signals, and underscored genetic and functional links between dietary intake and neural mechanisms. For example, the model identified a rare variant in rs76094283 (MAF = 0.0002 [imputation quality score =0.8]) as the most likely candidate to be the causal variant in the GDNF family receptor alpha 2 (*GFRA2*) locus. The variant achieved a PPA = 0.86 and showed suggestive evidence of association with increased fat intake with concomitant reduced carbohydrate and protein intake (*P*=4.3×10^−7^). The identified variant lies in the transcription start site of the *GFRA2* gene, which encodes the neurturin receptor implicated in neuron differentiation and survival.^38,39^ GFRalpha2-knockout mice display significant impairments in the parasympathetic and enteric nervous system, reduction in body weight, and memory of taste aversions, highlighting the importance of this molecular complex for eating behavior and nervous system development and function.^40,41^

**Fig 3.**
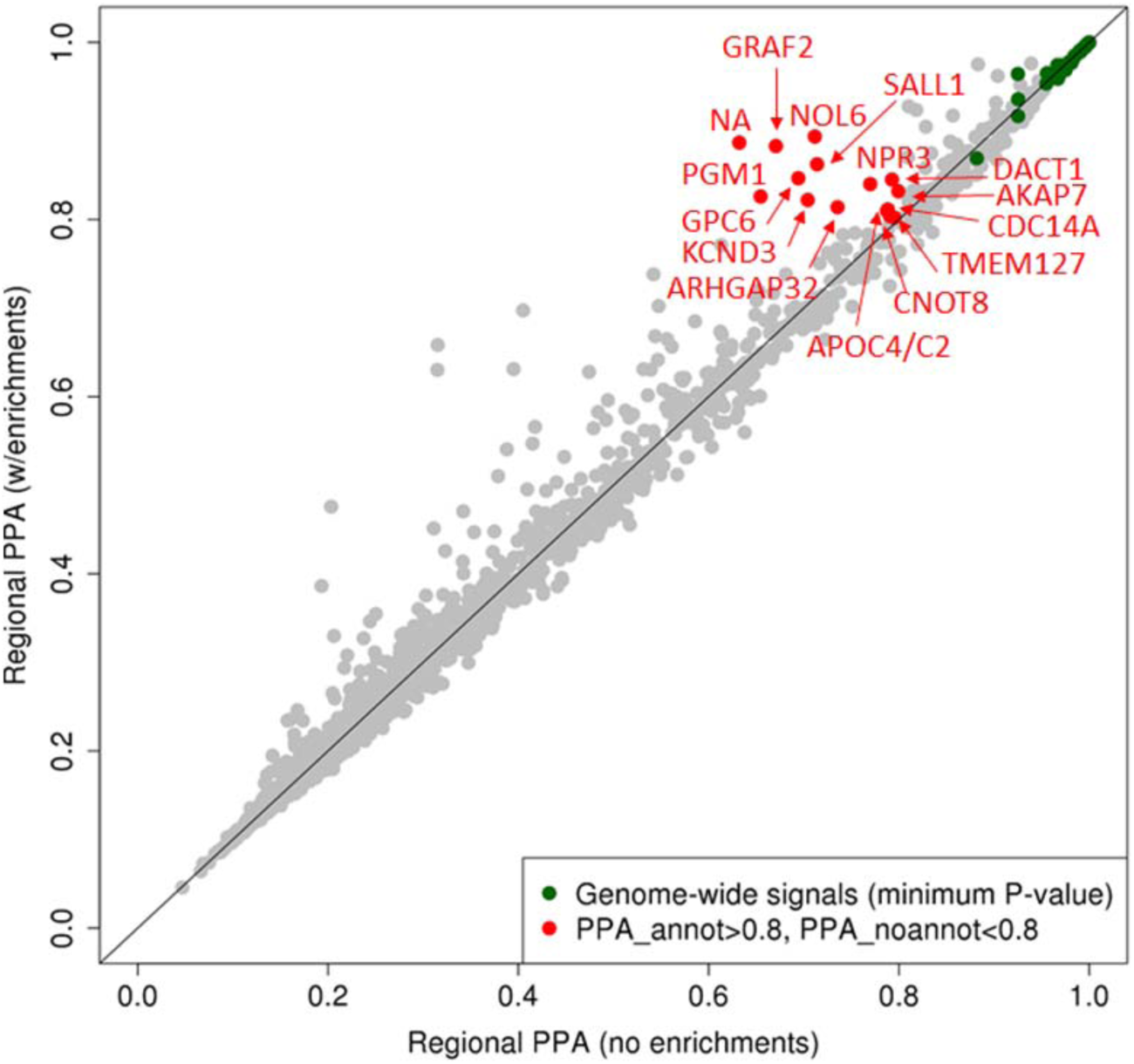
Joint analysis of functional genomic annotations and multi-trait genome-wide association meta-analysis. Reweighted multi-trait genome-wide association meta-analysis for dietary intake using fGWAS. Single significant annotations were combined using a hierarchical modeling approach based on penalized likelihood (Methods). The model identified DNase I hypersensitivity sites in fetal brain, weak enhancers in HeLa cells, and repressed chromatin in B cell as the most relevant annotations (Supplementary Table 14). Each point represents a region of the genome, and shown are the posterior probabilities of association (PPAs) of the regions in the models with and without the annotations.

To group the identified genome-wide significant loci into clusters with potential clinical similarities, we implemented a Bayesian nonnegative matrix factorization (bNMF) clustering algorithm (Methods).^42–44^ We aligned variants by their alleles associated with increased proportion of fat intake and their respective associations with 22 other dietary traits (Methods, Supplementary Fig. 3, Supplementary Table 21). The defining features of each identified cluster were determined by the most highly associated traits and variants after running 1,000 iterations. In 65% of the iterations, the data converged onto three clusters denoting different dietary composition patterns. In addition to increased proportion of fat intake in all clusters, cluster 1 was characterized by increased energy intake (“hyper caloric”), cluster 2 was driven by reduced proportion of protein and carbohydrate intake (“reduced carbohydrate and protein diet”), and cluster 3 was determined by increased proportion of protein intake (“increased protein diet”). Additional traits and defining loci for each cluster are described in Supplementary Fig. 4 and Supplementary Tables 22-23. In exploratory analyses, when aligning dietary intake variants for increased proportion of carbohydrate intake instead, clusters retained similar overall patterns.

We used the set of strongest-weighted variants from each cluster to generate partitioned polygenic scores (PPSs) (Methods). We first confirmed that PPSs were significantly associated with traits defining each cluster (Methods, Supplementary Fig. 3, Supplementary Table 24). We next assessed PPSs associations with BMI, T2D, and CAD in the UK Biobank. In the UK Biobank, the PPSs for clusters 2 (“reduced carbohydrate and protein diet”) and 3 (“increased protein diet”) were each associated with lower BMI with adjusted estimated effect sizes of −0.03 kg/m^2^ (SE=0.01, *P*=6.9×10^−8^) and −0.03 kg/m^2^ (SE=0.01, *P*=1.2×10^−16^) per allele increase in the PPS, respectively (Fig. 4). The PPS for the “increased protein diet” was associated with lower T2D with an adjusted estimated odds ratio of 0.97 (95%CI: 0.96 to 0.99; *P*=4.0×10^−4^) per allele increase in the PPS. We detected no association with CAD. To replicate findings from the UK Biobank, we further used data from the Partners Healthcare Biobank (*n*=19,563), an independent hospital-based biobank (Methods).^45^ In the Partners Healthcare Biobank, we noted a similar nominal association between the “reduced carbohydrate and protein diet” cluster and obesity (0.97, 95%CI: 0.95 to 0.99; *P*=0.02) and between the “increased protein diet” cluster and T2D (0.95, 95%CI: 0.91 to 0.99; *P*=0.01). Overall, these findings suggest that dietary patterns characterized by increased fat in combination with reduced carbohydrate intake may be associated with lower BMI, and that the combination of increased fat and protein intake may associated with lower T2D risk, but residual confounding may still exist.

**Fig 4.**
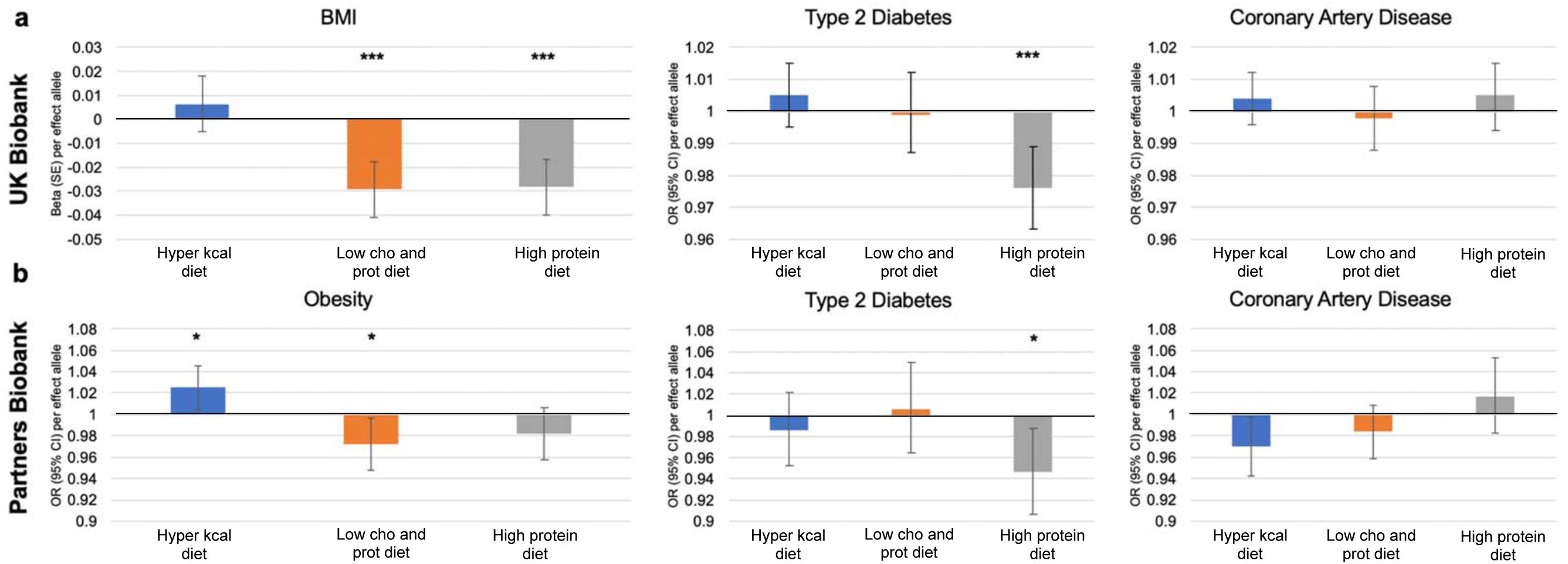
Association of cluster-specific polygenic risk scores and cardiometabolic phenotypes. Cluster-specific polygenic risk scores were defined by the top set of strongest-weighted variants for each cluster using a cutoff weighting of 1.09 (Methods). Individual participant scores were created by summing the number of alleles at each genetic variant weighted by the respective effect sizes on cluster pertinence. a). Individual-level data from the UK Biobank was used to test the association of cluster specific polygenic risk scores with BMI n=453,111), T2D (n=16,890 cases and 436,226 controls), and CAD (n=28,419 cases and 424,694 controls) after adjusting for age, sex, principal components, genotyping array, and BMI (only in analyses for CAD and T2D) (Methods). b) Individual-level data from the Partners Healthcare Biobank was used to investigate the extent to which previous observed associations replicated in an independent dataset. Sample size in Partners Healthcare Biobank was obesity (n=5,039 cases and 14,557 controls), T2D (n=1,497 cases and 18,099 controls), and CAD (n=2,673 cases and 16,293 controls) (Methods). Estimated effect sizes provided by one allele increase in the PRS. PRS associations were considered significant at Bonferroni corrected threshold of significance (0.05/12 (3 cluster and 3 outcomes) P<5.56×10^−3^). * P<0.05, ** P<0.01, ***P<<5.56×10^−3^

In the present study, we have expanded the genetic landscape of dietary intake by increasing the number of genome-wide significant loci from three to 96. Our findings provide insights into biological mechanisms that influence dietary intake, highlighting the relevance of brain-expressed genes, brain cell types, and central nervous system processes. Our results align with findings from tissue expression and gene-set enrichment analyses of BMI-associated loci that pinpoint the central nervous system as a critical regulator of energy homeostasis and body weight.^46,47^ A novel finding from this study is the enrichment of dietary intake associated signals for genes expressed in β1-tanycytes. Morphological studies have mapped β1-tanycytes on the floor and ventral parts of the third ventricle in the median eminence complex of the hypothalamus.^32,48^ Animal studies suggest that β1-tanycytes are nutrient and metabolite sensors that impact upon the blood-hypothalamus barrier plasticity and neuronal function.^34,49,50^ In this context, we also showed that β1-tanycytes are enriched for expression of genes previously associated with lipid homeostasis and growth factor, thus providing a strong cellular candidate for future studies. The multi-trait approach allowed us to leverage the observed genetic correlations among nutrients to increase power for locus discovery, as has been the case for other correlated traits.^51–53^ The bNMF clustering algorithm enabled us to dissect dietary intake genetic heterogeneity and identify three main domains of genetic variants that may have distinct impacts on obesity and T2D. Our findings, in line with previous evidence,^54^ may suggest that increasing proportion of fat intake in place of protein and carbohydrate associates with lower BMI. In addition, the combination of increased proportion of fat and protein intake associates with lower BMI and lower T2D risk. These findings are likely unaffected by the possibility that individuals at high BMI, T2D, or CAD genetic risk may have made changes in their diet, yet considering the complex network of biological and non-biological determinants of dietary intake, caution must be exercised in drawing strong conclusions.

Because the present study was limited to self-reported dietary intake, more precise measurements, including evaluating age and social, cultural, and economic factors as mediators or effect modifiers of a dynamic phenotype such as dietary intake may help refine our findings. In addition, expanding these analyses to non-European ancestries is warranted to determine the generalizability of the identified signals. Nonetheless, the present findings provide a starting point for understanding the biological variability of a complex and disease-relevant behavior, and provide specific supportive evidence for functional research that will aid in the discovery of mechanisms by which associated genes may affect dietary intake and related cardiometabolic risk.

## URLs

UK Biobank, http://www.ukbiobank.ac.uk/; CHARGE Consortium,; BOLT-LMM, https://data.broadinstitute.org/alkesgroup/BOLT-LMM/; LocusZoom, https://github.com/statgen/locuszoom-standalone/; Single-cell expression data, https://www.ncbi.nlm.nih.gov/geo/query/acc.cgi?acc=GSE93374; MAGMA software, http://ctg.cncr.nl/software/magma; MSigDB, http://software.broadinstitute.org/gsea/msigdb/collections.jsp; METAL software, http://genome.sph.umich.edu/wiki/METAL_Program; LDSC, https://github.com/bulik/ldsc; FUMA software, http://fuma.ctglab.nl/; GTEx, https://www.gtexportal.org/home/; DEPICT, https://data.broadinstitute.org/mpg/depict/; fGWAS, https://github.com/joepickrell/fgwas/; MACH, http://csg.sph.umich.edu//abecasis/MaCH/; SHAPEIT, https://mathgen.stats.ox.ac.uk/genetics_software/shapeit/shapeit.html; DGI db, http://dgidb.org/;

## Acknowledgements and Funding

This study was designed and carried out by the CHARGE nutrition Working Group Consortium. Part of this work was conducted using the UK Biobank resource under application number 27892. This study was supported by funding from European Commission award H2020-MSCA-IF-2015-703787 to J.M. H.S.D. and R.S. are supported by NIH R01 DK105072 and DK107859. R.S. is also supported by NIH R01 DK102696 and MGH Research Scholar Fund. J.M.L. is supported by NIH grants F32 DK102323 and T32 HL007901. C.S., J.C.F. and J.D. were supported by NIH grant U01 DK078616. J.C.F. is supported by NIH grant K24 DK110550. J.C. was supported by the American Diabetes Association Pathway to Stop Diabetes award 1-18-INI-14. T.H.P. and P.V.T. acknowledge the Novo Nordisk Foundation (Grant numbers NNF16OC0021496). T.H.P. acknowledges the Lundbeck Foundation (Grant number R190-2014-3904). Institutional Review Board and/or oversight committees approved the study in each participating cohort and all participants provided written informed consent. A full list of acknowledgments appears in Supplementary Note.

## Competing Financial Interests statement

A.Y.C. is currently employed by Merck Sharp & Dohme Corp., a subsidiary of Merck & Co., Inc., Kenilworth, NJ, USA. M.K.R. reports receiving research funding from Novo Nordisk, consultancy fees from Novo Nordisk and Roche Diabetes Care, and modest owning of shares in GlaxoSmithKline. All remaining authors declare no competing interests.

## Author Contributions

J.M., H.S.D., A.Y.C., D.I.C., J.C.F., and R.S., conceived and designed the study. J.C.F., and R.S. oversaw the study. H.S.D., J.M., C.S., J.M.L. and Y.S. served as analysts. Phenotype definitions were developed by J.M., H.S.D., J.M., and T.T. performed quality control and meta-analyses. Heritability and genetic correlation was performed by H.S.D. Bioinformatic analyses were performed and interpreted by J.M., H.S.D., C.S., J.M.L., Y.S., H.W., C.T., T.T., D.I.C., J.C.F., and R.S. Single-cell expression analyses were conducted and interpreted by J.M., P.V.T., T.H.P., J.C., L.T., and J.C.F. fGWAS was performed by C.S. and J.D. The bNMF clustering algorithm was developed and executed by M.S.U., and J.K. Figures were created by J.M., H.S.D., C.S., M.S.U., Y.S., P.V.T., T.H.P. J.D., D.I.C., and M.K.R. provided helpful advice and feedback on study design and the manuscript. J.M., H.S.D., C.S., J.C.F., and R.S., made major contributions to the writing and editing. All authors contributed to and critically reviewed the manuscript. All authors approved the final version of the manuscript.

## Data Availability

Summary GWAS statistics will be publicly available at the UK Biobank website (http://biobank.ctsu.ox.ac.uk/).

## Online Methods

### Samples

#### UK Biobank

The UK Biobank is a large population-based study established primarily to allow detailed investigations of the genetic and lifestyle determinants of a wide range of phenotypes.^15^ Data from >500,000 participants living in the United Kingdom who were aged 40-69 and living <25 miles from a study center were invited to participate between 2006-2010. We used data released in July 2017, and filtering (described below) resulted in a final sample size of 192,005 participants of European ancestry with dietary intake data. The UK Biobank received ethical approval from the National Research Ethics Service Committee North West Haydock (reference 11/NW/0382), and all study procedures were performed in accordance with the World Medical Association for medical research. The current study was conducted under UK Biobank application 27892.

#### CHARGE Consortium

The CHARGE Consortium was formed to facilitate GWAS among multiple large cohort studies.^55^ We used summary statistics of dietary composition from the CHARGE dietary composition GWAS (CHARGE; see URLs (to be added when available)) on 24 discovery cohorts which included up to 91,114 participants of European ancestry.^14^ All included studies were approved by local ethic committees, and informed consent was obtained from all the participants.

### Dietary intake assessment

#### UK Biobank

In the UK Biobank, dietary data were collected from 211,036 participants using the Oxford WebQ, a web-based 24-hr diet recall that asks participants to self-report on the frequency of intake of about 200 commonly consumed foods and drinks from the preceding 24 hours.^16,17^ The recall was administered at baseline visits in-person at assessment centers (towards the end of recruitment for the last 70,000 participants), and later administered electronically by email on four separate occasions over a 16-month period (Feb 2011 – April 2012) to ~320,000 participants with known email addresses. Email invitations were sent on variable days of the week in order to capture variability in daily dietary intake. Daily nutrient composition was estimated by multiplying the quantity consumed of each food or drink by the known nutrient composition as derived from the UK food composition database McCance and Widdowson’s The Composition of Foods and its supplements.^56^ The following analysis was limited to total energy (f.100002) and macronutrients intake (in grams per day; excluding supplements) of carbohydrate (f.100005), total fat (f.100004), and protein (f.100003). Recalls with improbable estimated total energy <500 or >4,000 kcal per day were excluded. For ~66% of the 211,036 participants who completed more than one recall, averages of each macronutrient were calculated as suggested elsewhere.^57^

#### CHARGE Consortium

As described in the CHARGE Consortium GWAS publication, assessment tools to estimate habitual dietary intake in the participating cohorts including validated cohort-specific food frequency questionnaires, diet history and diet records. Based on the responses to each dietary assessment tool and study-specific nutrient databases, habitual nutrient consumption was estimated. Over-reporters and under-reporters were excluded by standard cut-offs determined by each study cohort as part of quality control.^12^

### Genotyping and quality control

#### UK Biobank

We used genotype data released by the UK Biobank in July 2017. The genotype data collection, quality control, and imputation procedures are described elsewhere.^58^ In short, 489,212 participants were genotyped on two customized genetic arrays (the UK BiLEVE Axiom array [*n* = 50,520] and the UK Biobank Axiom array [*n* = 438,692]) covering 812,428 unique genetic markers (95% overlap in variant content). Quality control procedures were conducted by the UK Biobank. Samples were removed for high missingness or heterozygosity (968 samples) and sex chromosome abnormalities (652 samples). Variants were tested for batch effects (197 variants/batch), plate effects (284 variants/batch), Hardy-Weinberg equilibrium (572 variants/batch), sex effects (45 variants/batch), array effects (5,417 genetic variants), and discordance across control replicates (622 on UK BiLEVE Axiom array and 632 UK Biobank Axiom array) (*P*<10^−12^ or <95% for all tests). For each batch (106 batches total) markers that failed at least one test were set to missing. Before imputation, 488,377 individuals and 805,426 SNPs pass QC in at least one batch (>99% of the array content). Genotypes were phased and imputed by the coordinating team to approximately 96 million genetic variants by using a combined reference panel, including the Haplotype Reference Consortium and the UK10K haplotype panel. Imputed and quality-controlled genotype data were available for 487,422 individuals and 92,693,895 genetic variants. Population structure was captured by principal component analysis on the samples using a subset of high quality (missingness <1.5%), high frequency variants (>2.5%) (~100,000 genetic variants) and identified the sub-sample of white British descent. We further clustered participants into four ancestry clusters using K-means clustering on the principal components, identifying 453,964 participants of European ancestry included in the present analysis, of which 192,005 had available dietary intake data and thus remained in the final analysis.

#### CHARGE Consortium

Genotyping and quality control methods for the CHARGE Consortium dietary intake GWAS has been detailed elsewhere.^14^ In brief, each participating cohort performed quality control for genotyped variants based on minor allele frequency (MAF), call rate, and departure from Hardy-Weinberg Equilibrium. Phased haplotypes from 1000G were used to impute ~38 million autosomal variants using a Hidden Markov Model algorithm implemented in MACH/minimac^59,60^ or SHAPEIT/IMPUTE.^61,62^ Variants with low minor allele count (MAC<20) in the meta-analysis and low imputation quality (<0.4) were removed. The number of autosomal genetic variants analyzed in this study was ~11.8 million.

### Single-trait and multi-trait genome-wide association meta-analysis

In the UK Biobank, genetic association analyses were performed separately for carbohydrate, fat, and protein as percentages of total energy in 192,005 participants using BOLT-LMM linear mixed models and an additive genetic model adjusted for age, sex, 10 principal components of ancestry, genotyping array and genetic correlation matrix [jl2] with a maximum per SNP missingness of 10% and per sample missingness of 40%.^19^ We used a minor allele frequency threshold of 0.001. In a second model, BMI was added to the covariates to account for genetic effects mediated through body composition.

To maximize the statistical power, we meta-analyzed the dietary intake GWAS in UK Biobank and the CHARGE Consortium. Meta-analysis was performed using METAL by weighting effect size estimates using the inverse of the corresponding standard errors squared (version – released March 25 2011).^63^ The genetic signals correlated strongly between the two samples supporting the meta-analysis genome-wide (*r_g_*>0.6 all *P* <0.001; Supplementary Table 2). We assessed heterogeneity in genetic effects across studies using the *I*^2^ heterogeneity index.^64^

Single-trait estimates for carbohydrate, fat, and protein from the meta-analyses were combined in a multi-trait analysis using the cross-phenotype association software (CPASSOC).^65^ In brief, CPASSOC uses summary-level data from single variant-trait associations from GWAS to boost the statistical power for locus discovery by leveraging the observed genetic correlation between traits. The joint analysis of multiple phenotypes using CPASSOC provides two statistics, S_Hom_ and S_Het_. S_Hom_ is similar to statistics generated by the fixed-effects meta-analysis method, but uses the sample size for a trait as a weight instead of the variance and accounts for the correlation of summary statistics among traits and studies induced by correlated traits, potential overlapping or related samples. S_Het_ is an extension of S_Hom_ that improves power when the genetic effect sizes vary for different traits. The distribution of S_Het_ values under the null hypothesis of no association was obtained through an estimated beta distribution. In a recent comparison of statistical power of 19 multivariate genome-wide association methods, it was shown that S_Het_ seems to benefit from the presence of opposite effect estimates, which is relevant for dietary composition. In addition, Shet performed slightly better when applied to highly correlated traits.^66^

### Heritability and genetic correlation

Trait SNP-based heritability was calculated as the proportion of trait variance due to additive genetic factors using BOLT-REML.^19^ Genomic control lambda (λ_GC_) values were calculated using GenABEL in R^67^ using post-quality control GWAS results. LD Score regression was used to estimate the cross-cohort genetic correlations.^18^ Calculated linkage disequilibrium (LD) scores from 1000G European reference population were obtained online (see URLs).

### Genomic risk loci definition

Distinct associated loci from the meta-analysis were defined using the Functional Mapping and Annotation FUMA platform^68^ (see URLs). We first defined distinct significant genetic variants, which had a genome-wide significant *P*-value (*P*<5×10^−8^) and were in low LD (*r^2^*<0.6). These variants were further represented by lead genetic variants, which are a subset of the distinct significant variants that are in approximate LD with each other at *r^2^*<0.1 (based on LD information from UK Biobank genotype data). Subsequently, genomic risk loci were defined by merging lead variants that physically overlapped or for which LD blocks were less than 250 kb apart. Borders of the associated genomic loci were defined by identifying all variants in LD (*r^2^*≥0.6) with one of the distinct significant variants in the locus, and a region containing all of these candidate variants was considered to be a single distinct genomic locus.

### Genetic risk for cardiometabolic diseases and dietary intake composition

To assess whether our findings were influenced by the possibility that individuals at high metabolic risk had made changes to their diet (reverse causation), we investigated the associations of polygenic risk scores (PRS) for BMI, T2D, and CAD with macronutrient intake. Individual-level data from the UK Biobank were used to create individual participant scores by summing the number of alleles at each genetic variant weighted by the respective effect sizes on BMI, T2D, and CAD based on estimated effect sizes from published meta-analyses of GWAS for BMI (n=339,224 individuals),^46^ T2D (n=74,124 T2D cases and 824,006 controls),^69^ and CAD (60,801 cases and 123,504 controls).^70^ We compared mean proportion of total energy intake from carbohydrate, fat, and protein among quartiles of each PRS using analysis of variance.

### Gene mapping and functional annotation

All genetic variants in the meta-analysis results that were in LD (*r^2^*>0.6) with one of the distinct significant variant and that had *P*<5×10^−8^ and MAF > 0.0001 were mapped to genes in FUMA using positional mapping, expression quantitative trait loci (eQTL) mapping, and chromatin interaction mapping. Predicted functional consequences for identified variants were obtained from databases containing known functional annotations, including ANNOVAR categories,^71^ Combined Annotation Dependent Depletion (CADD) scores,^72^ RegulomeDB scores,^73^ and chromatin states.^74,75^ ANNOVAR categories identify the variants’ genic position and associated function. CADD scores predict how deleterious the effect of a variant is likely to be for protein structure/function, with higher scores referring to higher deleteriousness. The RegulomeDB score is a categorical score based on information from eQTLs and chromatin marks, ranging from 1 to 7, with lower scores indicating an increased likelihood of having a regulatory function. The chromatin state represents the accessibility of genomic regions with 15 categorical states predicted by a hidden Markov model based on 5 chromatin marks for 127 epigenomes in the Roadmap Epigenomics Project.^75^ A lower state indicates higher accessibility, with states 1-7 referring to open chromatin states.

### Tissue and gene-set enrichment analyses

We used multi-trait meta-analysis *P*-values as input for an enrichment analysis in 53 tissue types obtained from the Gene-Tissue Expression Project (GTEx)^30^ and 10,651 predefined gene-sets derived from MSigDB^31^ in MAGMA.^76^ For tissue enrichment analyses, gene expression values are log2 transformed average RPKM per tissue type after winsorized at 50 based on GTEx RNA-seq data. MAGMA was performed using the result of gene analysis (gene-based *P*-value) and tested for one side (greater) with conditioning on average expression across all tissue types. For gene-set enrichment we tested for association in types of predefined gene sets representing known biological functions and metabolic pathways derived from MSigDB.

### Brain single-cell-expression analyses

To connect genomic loci implicated in dietary composition with the specific brain cell types defined by gene expression profiles, we used two independent brain single-cell RNA sequencing datasets. We first used information from 24 level 1 brain cell types (level 1 clusters were characterized based on expression of known marker genes) and 149 level 2 brain cell types (subtypes of a level 1 grouping (for example, medium spiny neurons expressing Drd1 or Drd2) from Skene et al.^33^ In brief, brain-cell-type expression data were drawn from single-cell RNA-seq data from mouse brains. For each gene, the value for each cell type was calculated by dividing the mean unique molecular identifier (UMI) counts for the given cell type by the summed mean UMI counts across all cell types. Single-cell gene sets were derived by grouping genes into 40 equal bins by specificity of expression. These gene sets were tested for association with gene-based test statistics using MAGMA.^76^

Given the relevance of the hypothalamic Arc-ME in energy homeostasis and appetite control,^10,32^ we used Drop-seq data from 50 transcriptionally distinct Arc-ME cell populations described in Campbell et al.^32^ Drop-seq data of mouse Arc-ME (GSE93374) were averaged using the Seurat Average Expression function. Two different levels of clustering were used: a coarser clustering across all identified cell types shown and a finer-grained clustering limited to neuronal cell types as described.^32^ To account for droplet-specific differences in mRNA captures rates, we log-normalized the averaged counts using the default Seurat “logNormalize” method. This divides each observed expression value by the sum for that gene, multiplies by a scaling factor (here the default, 10000), adds 1, and takes the natural log. Genes were mapped from mouse gene symbols to mouse Ensembl gene identifiers (using Ensembl version 83) and to the human Ensembl gene identifiers (using Ensembl version 82). Mouse gene identifiers mapping to several mouse Ensembl identifiers were discarded and the human gene with the highest degree of mouse homology was retained in instances where a mouse gene mapped to several human genes. Expression levels of mouse genes mapped to the same human gene were averaged. A two-step z-score procedure was applied such that the expression levels for each gene were transformed to standard normal distributions and the expression levels for each cell cluster were transformed to standard normal distributions. The resulting matrices were used to run DEPICT^77^ vs1.194 (https://github.com/perslab/depict/) to prioritize cell clusters on the basis of summary statistics for multi-trait, carbohydrate, protein, fat intake (GWAS association *P*-value cutoff <10^−5^). DEPICT cell type enrichment *P*-values were adjusted for the number of cell types tested for enrichment for each trait, but not for the overall number of traits tested. Code to construct the average cell type expression and the DEPICT configuration file can be found at https://github.com/perslab/dietpref_merino.

### Joint analysis of functional genomic annotations

We used fGWAS,^37^ a hierarchical modeling approach that incorporates multiple types of genetic annotations to re-weight association measures by using information from the most relevant annotations. The main sources of genomic annotation were elements of gene structures, outputs from genome segmentation of the six main ENCODE cell lines, and 13 maps of DNase-I hypersensitivity from primary fetal brain cell types and cell lines. Details of the generation of these annotations have been described elsewhere.^78^

We first tested each of the genomic annotations individually and estimated the degree of enrichment of each specific annotation with loci that influence dietary intake. The annotation with the most significant enrichment was retained and tested jointly with each remaining annotation. If the most significant two annotation model improved the model likelihood, then the two annotations in the model were retained and the process continued until the model likelihood cannot be improved. This process resulted in the “best joint model”. By default, fGWAS partitions the genome into “blocks” of 5,000 genetic variants. We used a modified approach by delineating 1Mb windows comprising all genetic variants within 500 kb of the index variant and partitioned the intervening regions into ~1Mb windows. For non-genome-wide significance regions, we used the default approach. These windows were given as input into fGWAS using the --bed command and a separate fGWAS analysis was performed using only the set of annotations remaining in the “best joint model”. The genome-wide enrichments were used as priors in a Bayesian fine-mapping analysis implemented in fGWAS to calculate posterior probabilities for each region and each genetic variant in the designated windows.

### Bayesian nonnegative matrix factorization (bNMF) algorithm

We applied the bNMF clustering algorithm^42–44^ with the aim of grouping identified dietary intake genetic loci into subgroups of variants based on potential similarities across diverse dietary intake traits. The main input for the bNMF algorithm was the set of the 96-dietary intake associated variants identified in this study. We restricted the analysis to the 79 variants that showed at least nominal significant association with fat intake. Given that C/G and A/T alleles are ambiguous and can lead to errors in aligning alleles across GWAS, we avoided inclusion of ambiguous alleles, choosing proxies instead. Next, publicly available summary association statistics for 22 dietary intake traits from the UK Biobank^79^ were aggregated for each dietary intake variant (Supplementary Table 21). For the 79-dietary intake associated variants, the fat intake– increasing alleles were identified and all future analyses used the aligned fat intake–increasing alleles. We aligned for fat intake as opposed to other macronutrients since we had the highest confidence in determining allele association directionality. We generated standardized effect sizes for variant trait associations from GWAS by dividing the estimated regression coefficient by the standard error, using the UK Biobank summary statistic results (variant-trait association matrix (96 by 22)). To enable an inference for latent overlapping modules or clusters embedded in variant-trait associations, we modified the existing bNMF algorithm to explicitly account for both positive and negative associations as was done previously.^43,44^ The defining features of each cluster were determined by the most highly associated traits, which is a natural output of the bNMF approach. bNMF algorithm was performed in R for 1,000 iterations with different initial conditions, and the maximum posterior solution at the most probable number of clusters was selected for downstream analysis.

### Polygenic risk scores for trait and outcome association with each cluster

The results of the bNMF algorithm provide cluster-specific weights for each variant and trait. Variants and traits defining each cluster were based on a cut-off of weighting of 1.09 which was determined by the optimal threshold to define the beginning of the long-tail of the distribution of cluster’s weights across all clusters (top 5% were considered to be significant). Polygenic risk scores (PRSs) were created by summing the number of alleles at each genetic variant weighted by the respective effect sizes on cluster pertinence. To confirm that cluster-specific PRSs mirrored traits defining each cluster, we tested for associations of PRS with each GWAS dietary trait using inverse-variance weighted fixed effects meta-analysis. A conservative significance threshold was set at 7.6×10^−4^ (0.05/66; 22 traits and 3 clusters).

Next, individual-level data from the UK Biobank were used to create individual participant scores by summing the number of alleles at each genetic variant weighted by the respective effect sizes on cluster pertinence. Scaling of the individual PRS was performed to allow interpretation of the effects as a per-1 risk allele increase in the PRS for each trait (division by twice the sum of the effect sizes and multiplication by twice the square of the SNP count representing the maximum number of risk alleles). We tested the association of each cluster-specific PRS with BMI (n=453,111), T2D (n=16,890 cases and 436,226 controls), and CAD (n=28,419 cases and 424,694 controls) adjusting for age, sex, 10 principal components, genotyping array, and BMI (only in analyses for CAD and T2D). We next used individual-level data from the Partners Healthcare Biobank (*n* =19,596)^45^ to investigate the extent to which UK Biobank associations replicated in an independent dataset. A detailed description of the Partners Healthcare Biobank is provided below. Disease prevalence for obesity (n=5,039 cases and 14,557 controls), T2D (n=1,497 cases and 18,099 controls), and CAD (n=2,673 cases and 16,293 controls) were determined from electronic medical records using both structured and unstructured data. Logistic regression models were adjusted for age, sex, 5 principal components, and obesity (only in analyses for CAD and T2D). PRS associations with cardiometabolic phenotypes were considered significant at Bonferroni corrected threshold of significance at *P<*5.6×10^−3^ (0.05/9; 3 cluster and 3 traits).

### Partners Healthcare Biobank

The Partners HealthCare Biobank maintains blood and DNA samples from more than 60,000 consented patients seen at Partners HealthCare hospitals.^45^ Patients are recruited in the context of clinical care appointments and also electronically through the patient portal at Partners HealthCare. Genomic data for 19,596 participants of European ancestry were generated with the Illumina Multi-Ethnic Genotyping Array. The genotyping data were harmonized, and quality controlled with a three-step protocol, including two stages of genetic variant removal and an intermediate stage of sample exclusion. The exclusion criteria for variants were: 1) missing call rate ≥0.05, 2) MAF <0.001, and 3) deviation from Hardy-Weinberg equilibrium (*P* < 10^−6^). The exclusion criteria for samples were: 1) sex discordances between the reported and genetically predicted sex, 2) missing call rates per sample ≥0.02, 3) subject relatedness (pairs with estimated identity-by-descent ≥0.125, from which we removed the individual with the highest proportion of missingness), and 4) population structure showing more than four standard deviations within the distribution of the study population, according to the first four principal components. Phasing was performed with SHAPEIT2^80^ and then then imputations were performed with the Haplotype Consortium Reference Panel^81^ using the Michigan Imputation Server.^82^ Written consent was provided by all study participants. Approval for analysis of Biobank data was obtained by Partners IRB, protocol # 2018P002276.

