## Supplemental Dataset for "Multi-trait genome-wide association meta-analysis of dietary intake identifies new loci and genetic and functional links with metabolic traits"

### Slide 1
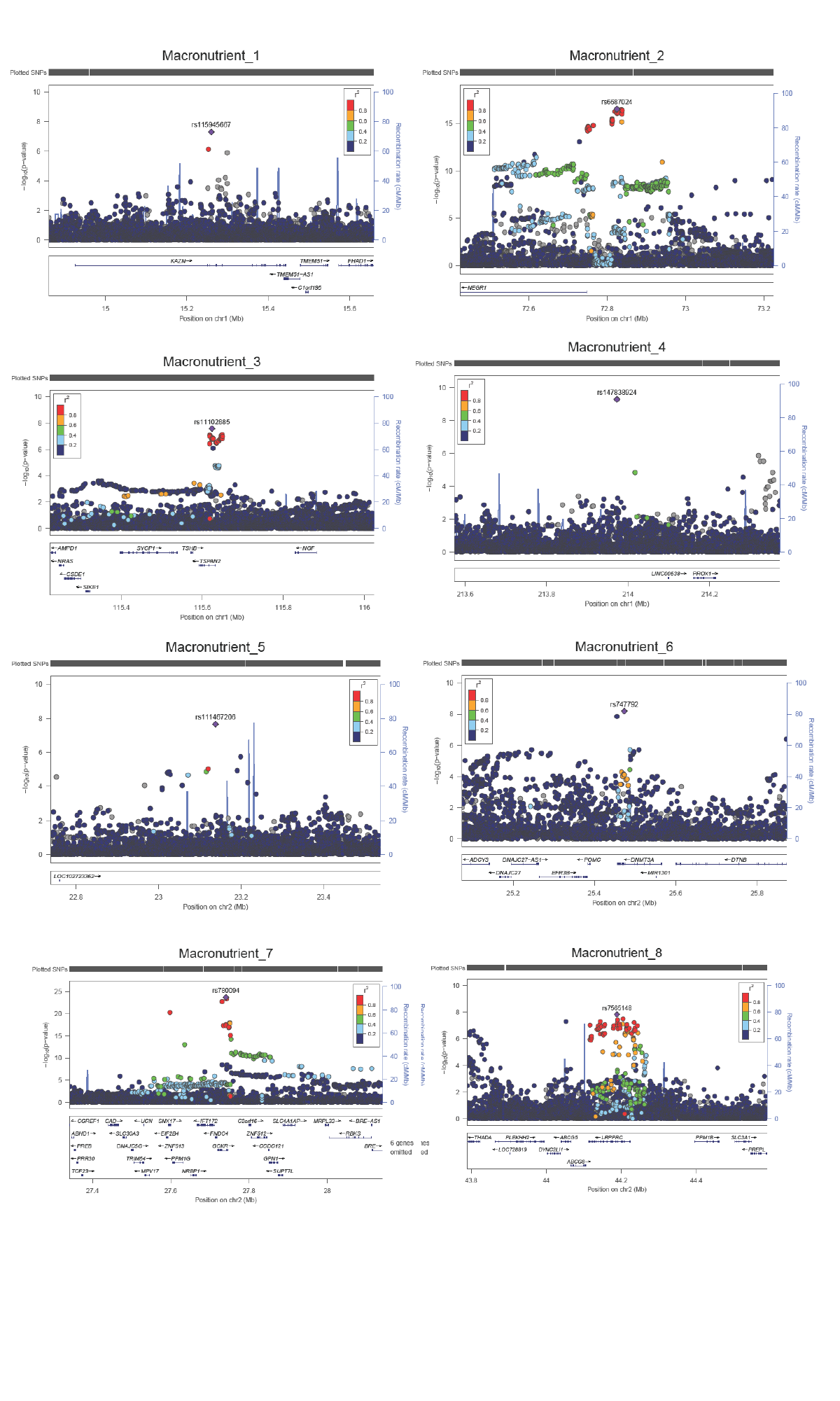

### Slide 2
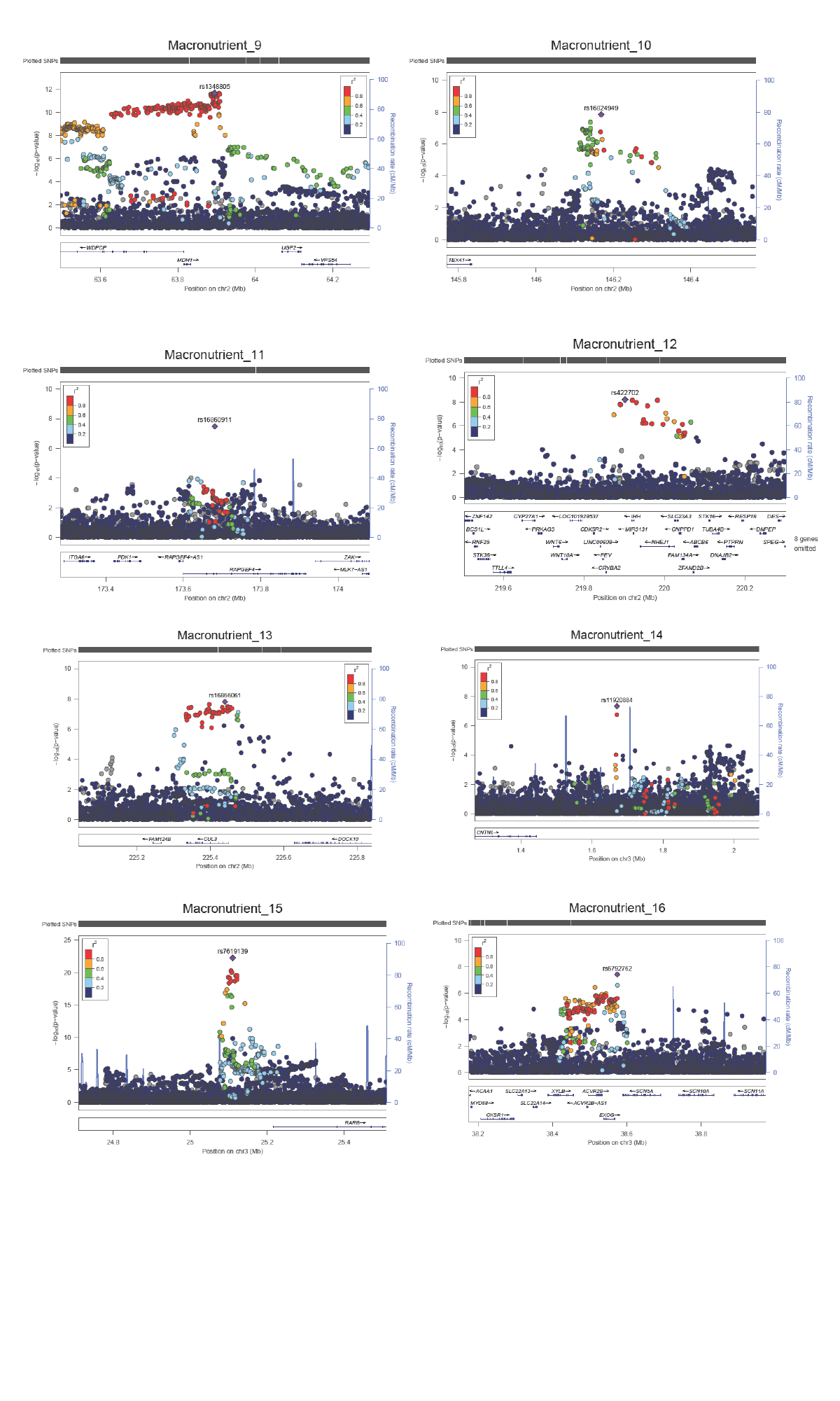

### Slide 3
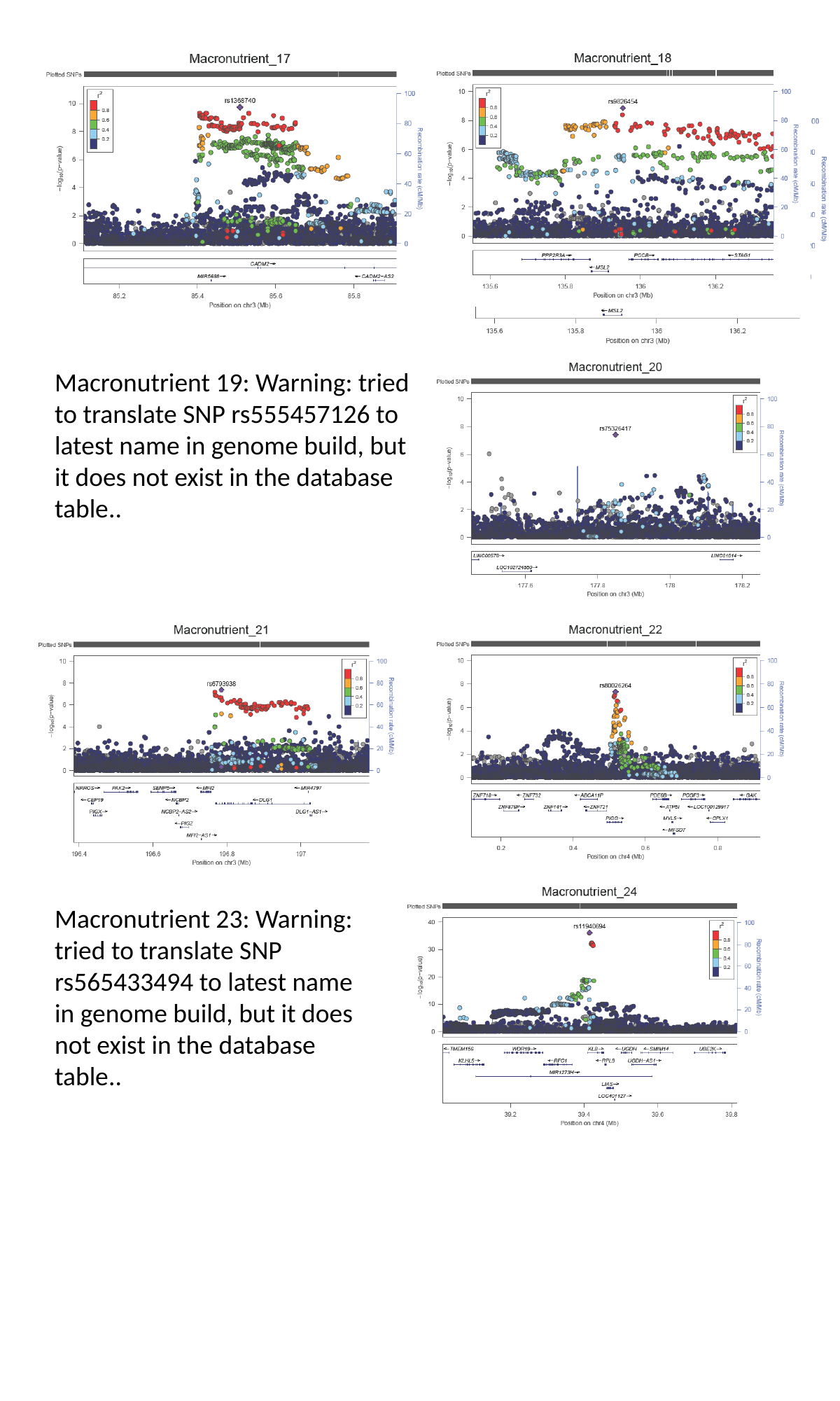

Macronutrient 19: Warning: tried to translate SNP rs555457126 to latest name in genome build, but it does not exist in the database table..
Macronutrient 23: Warning: tried to translate SNP rs565433494 to latest name in genome build, but it does not exist in the database table..

### Slide 4
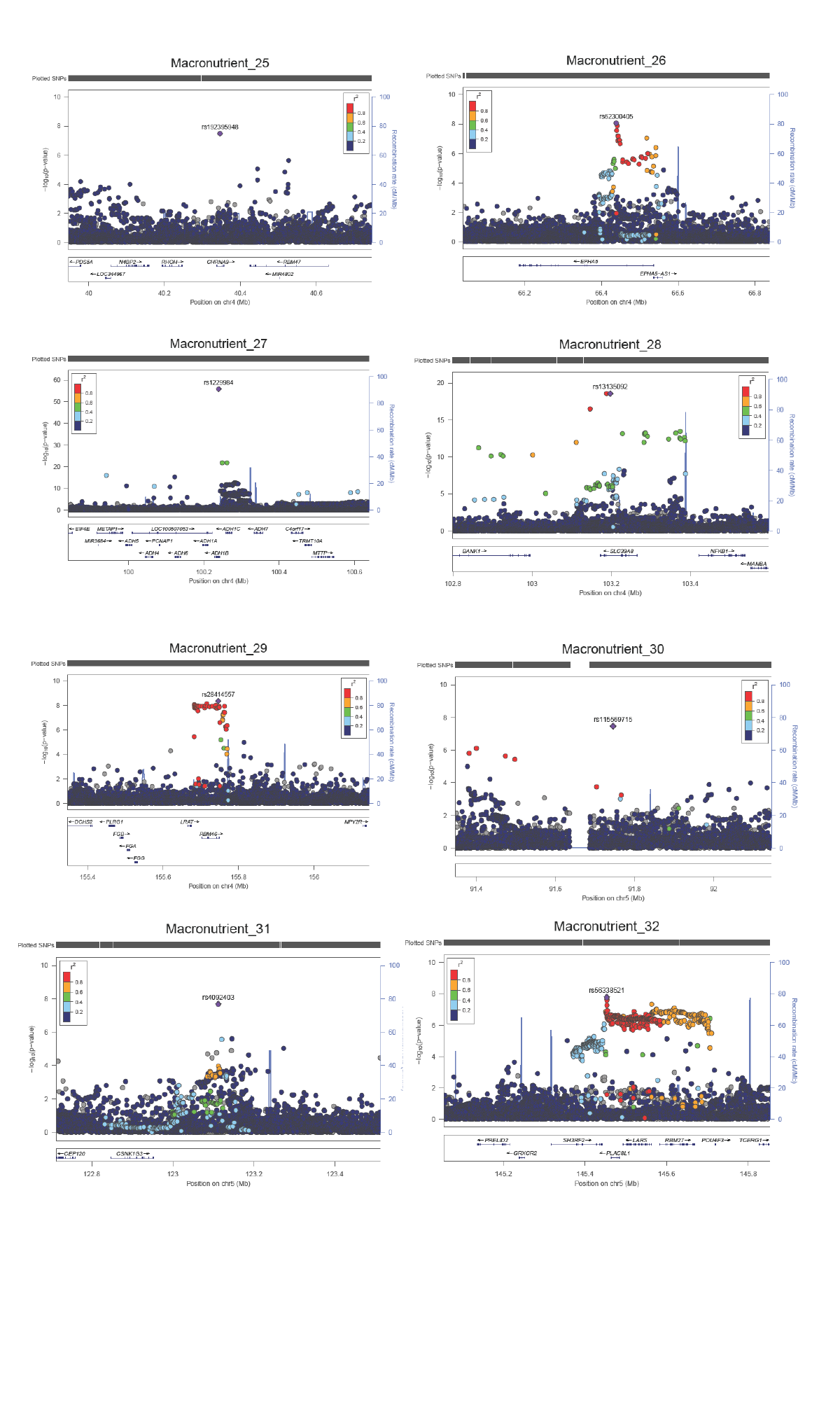

### Slide 5
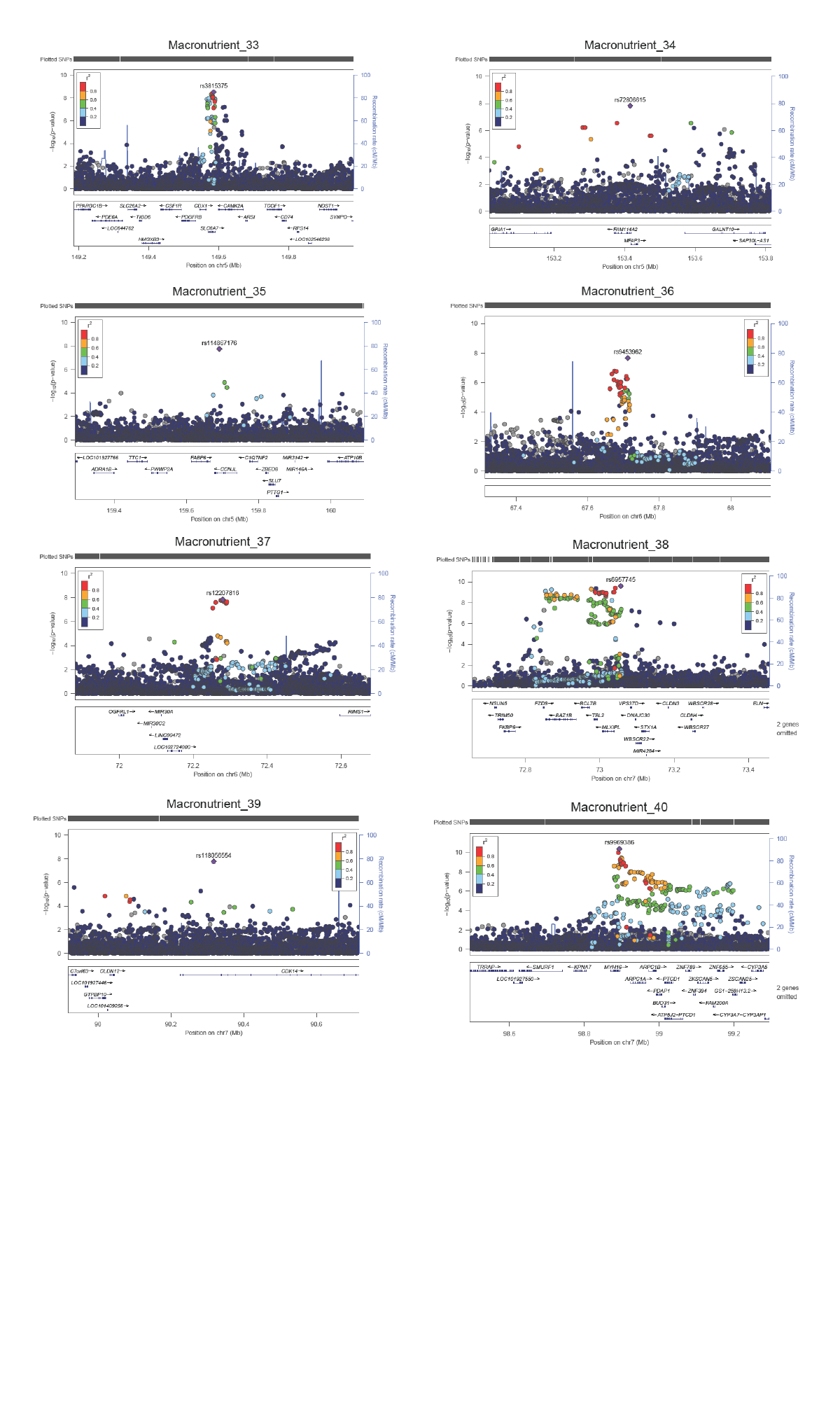

### Slide 6
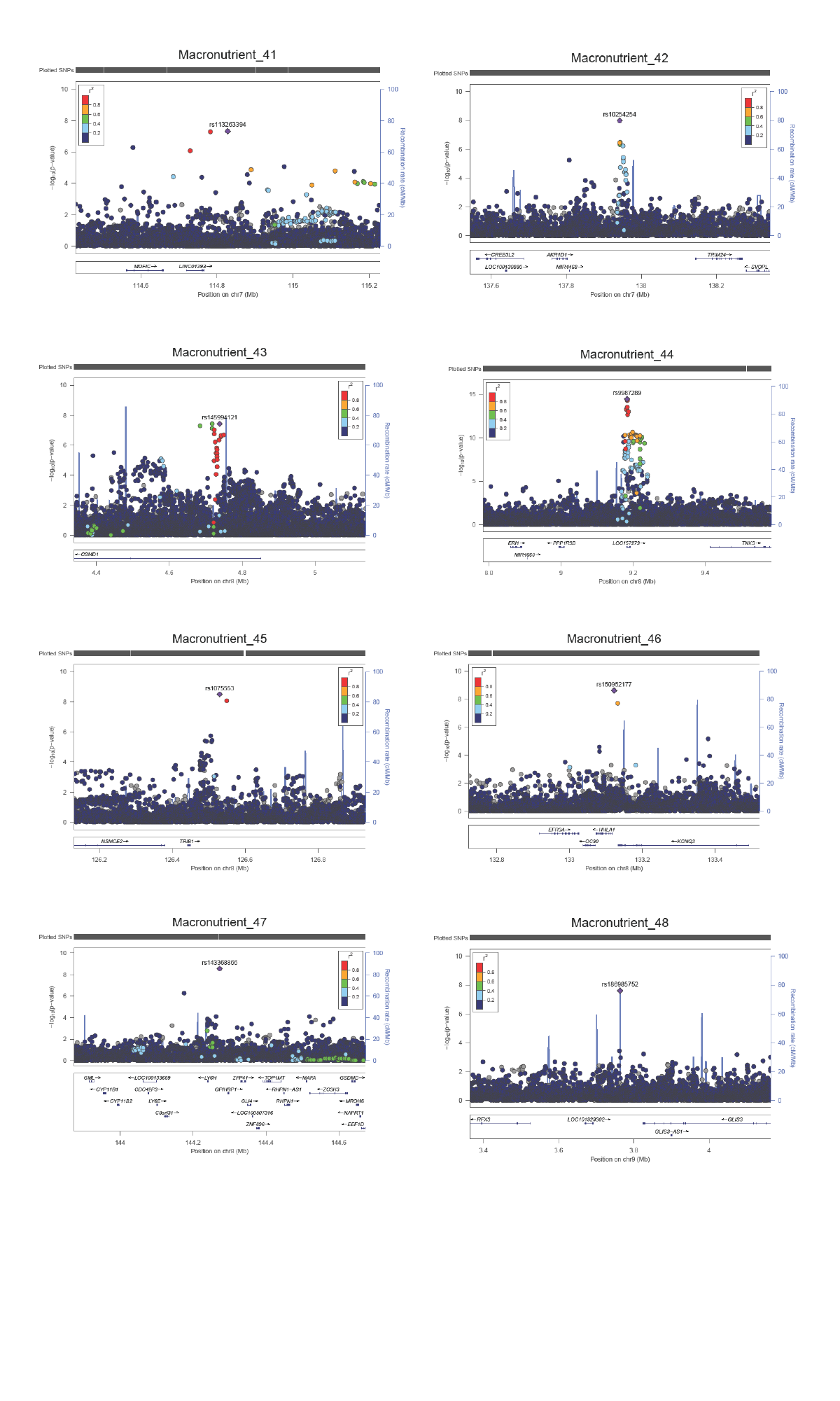

### Slide 7
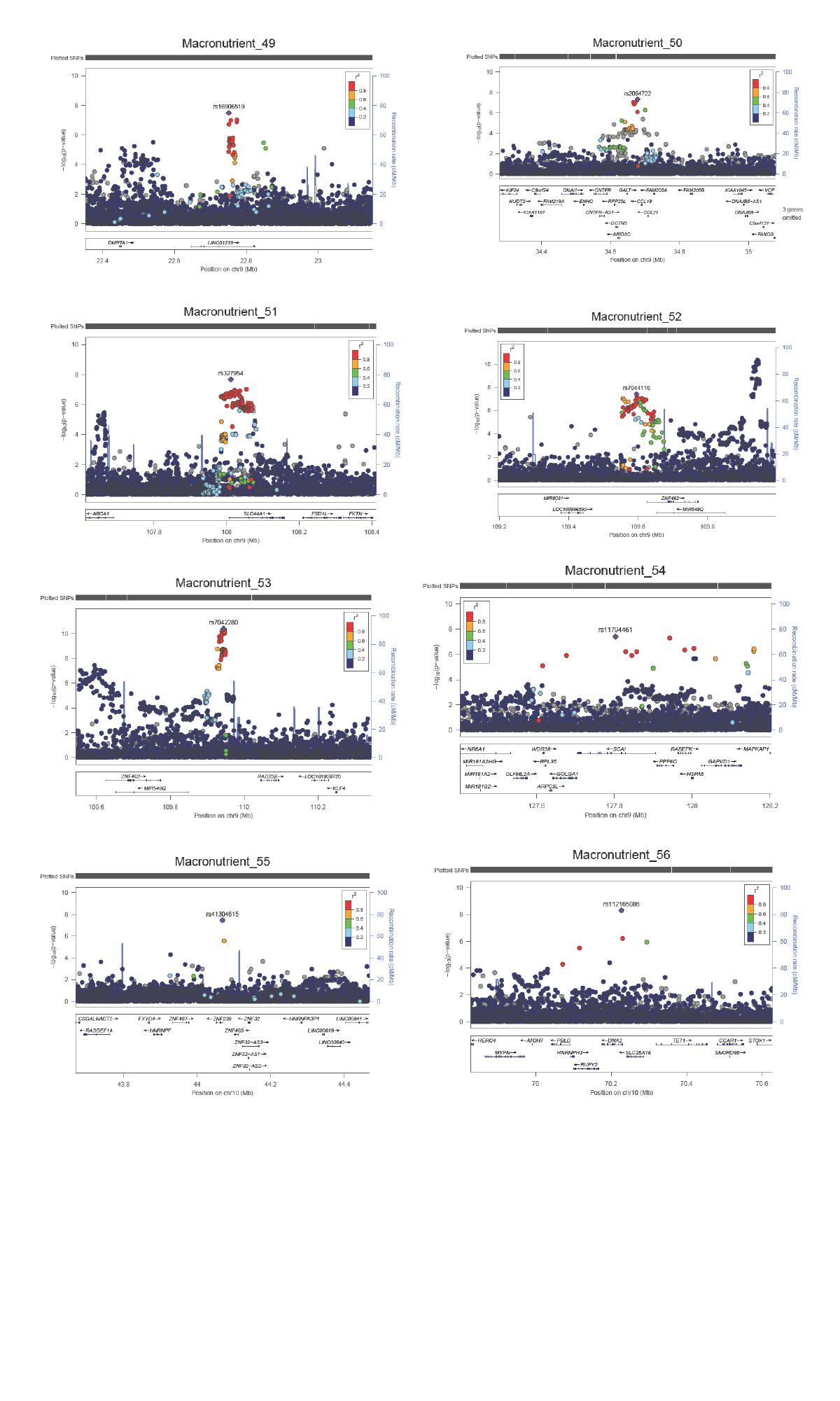

### Slide 8
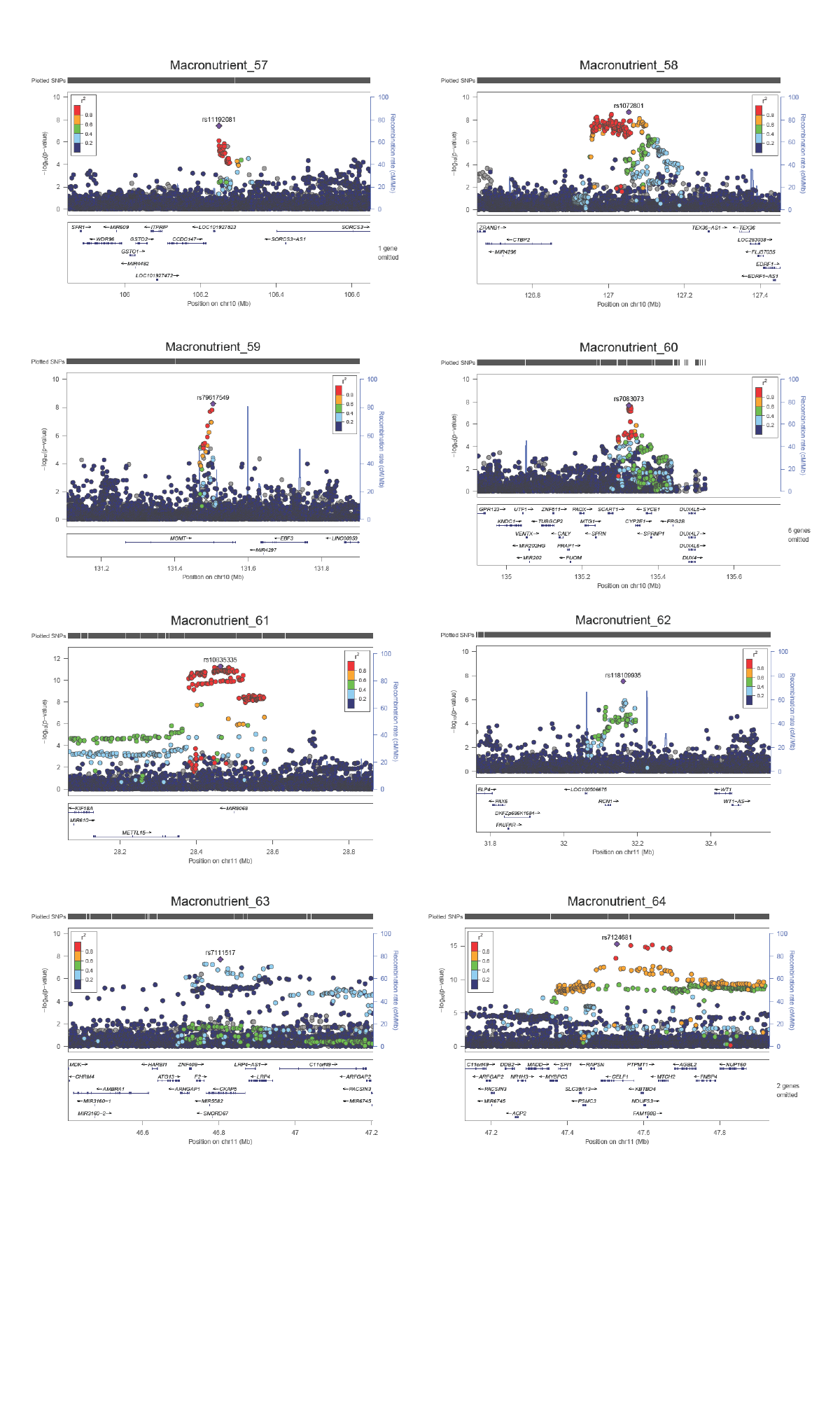

### Slide 9
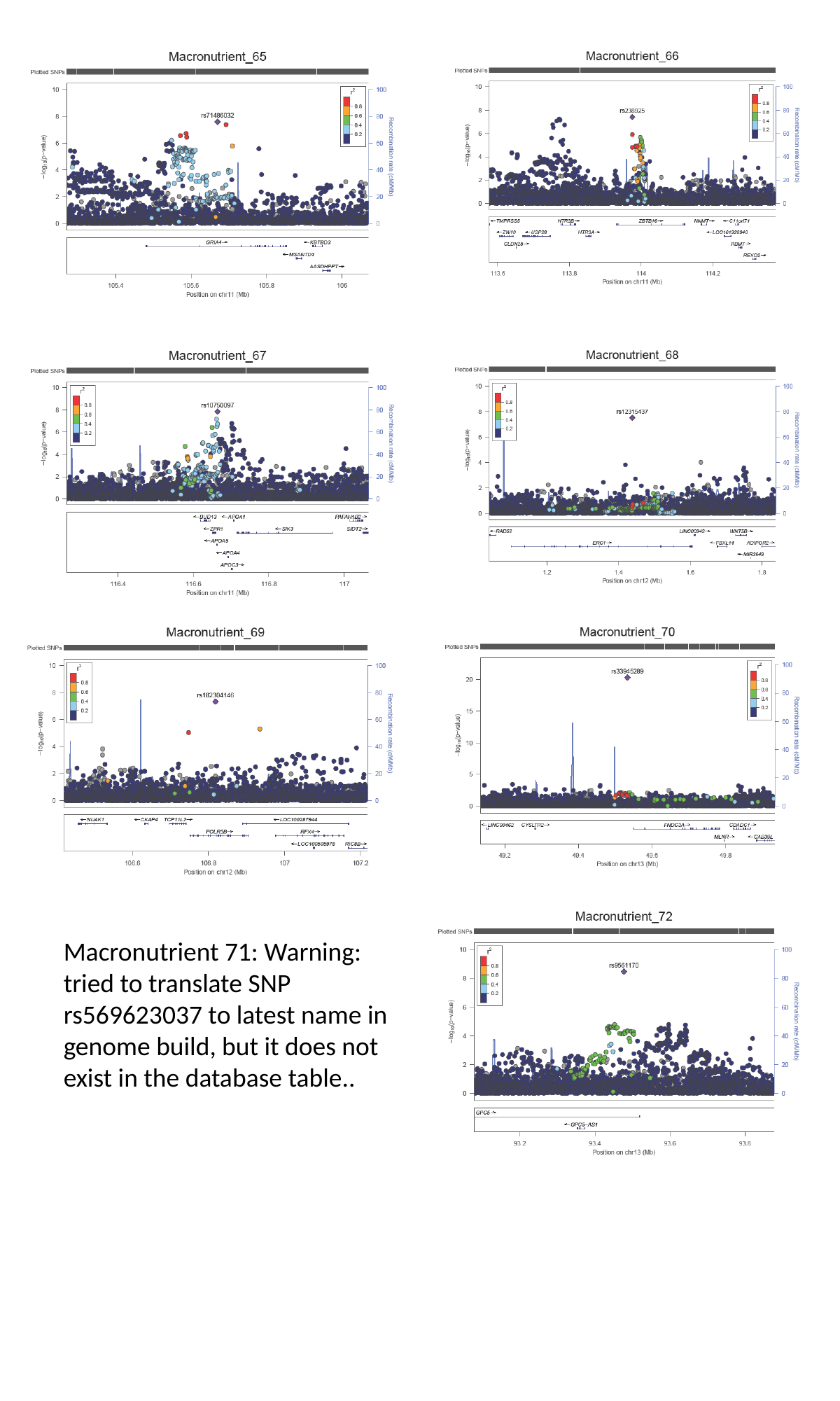

Macronutrient 71: Warning: tried to translate SNP rs569623037 to latest name in genome build, but it does not exist in the database table..

### Slide 10
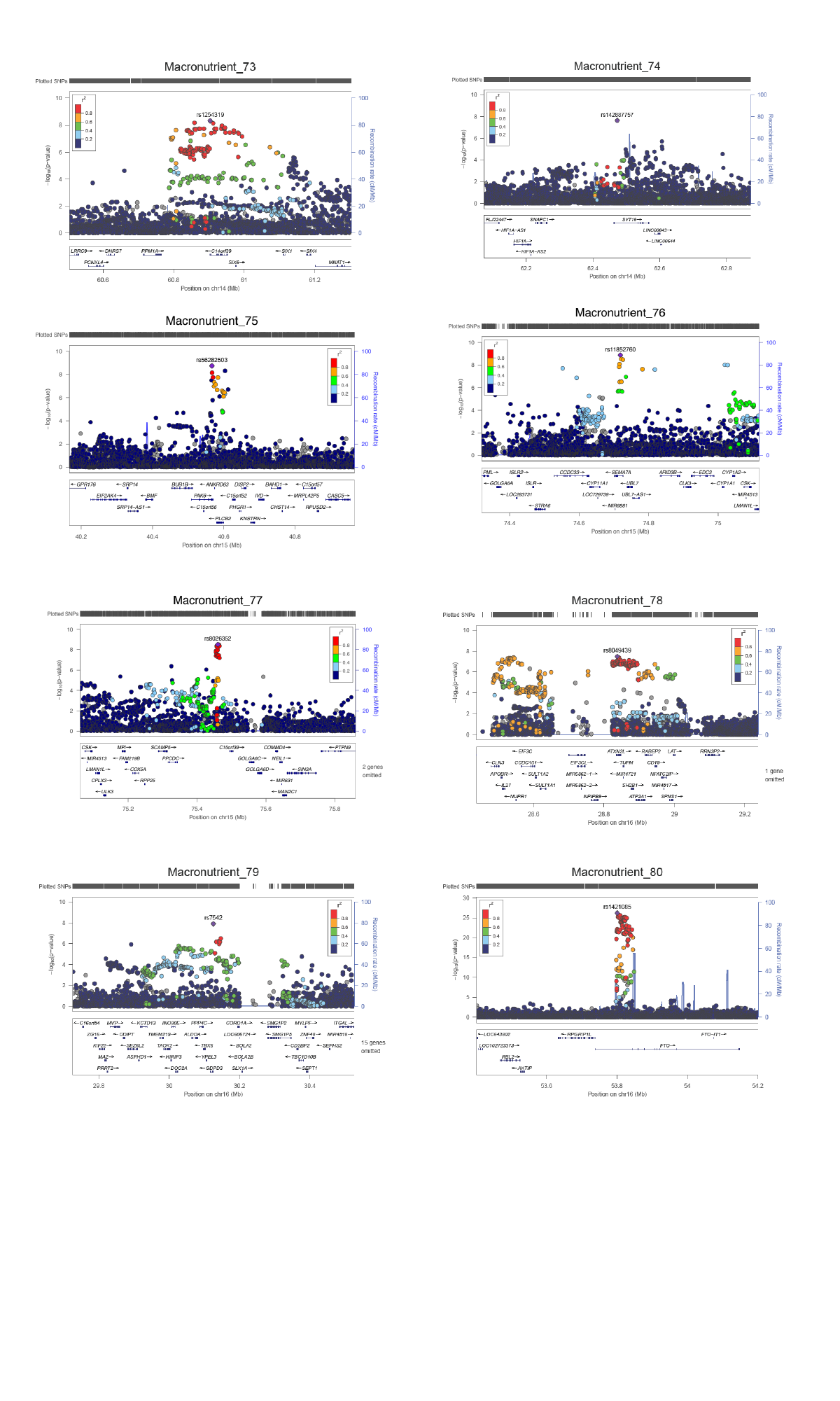

### Slide 11
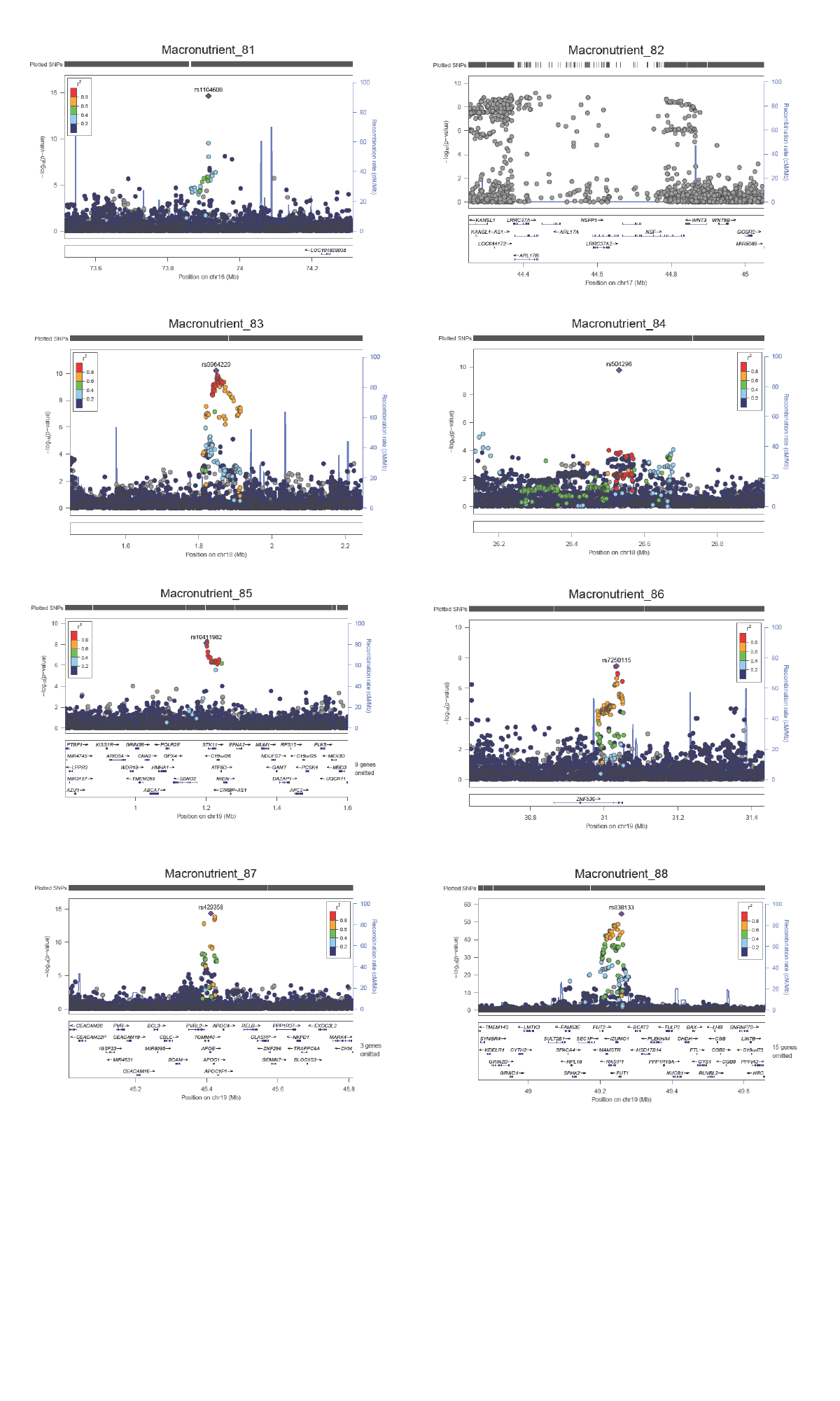

### Slide 12
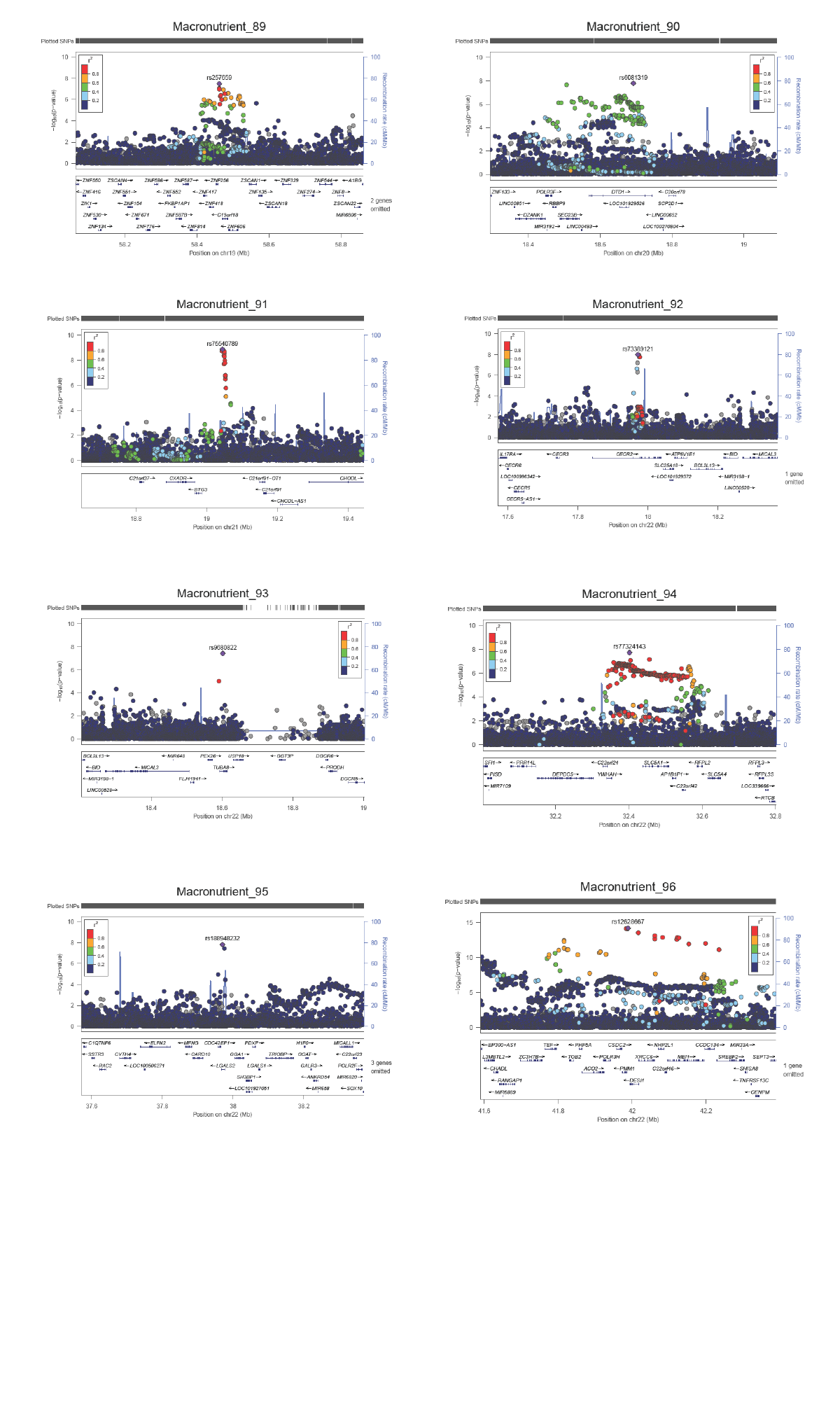
