## Supplemental Figures for "Multi-trait genome-wide association meta-analysis of dietary intake identifies new loci and genetic and functional links with metabolic traits"

**Supplementary Figures**

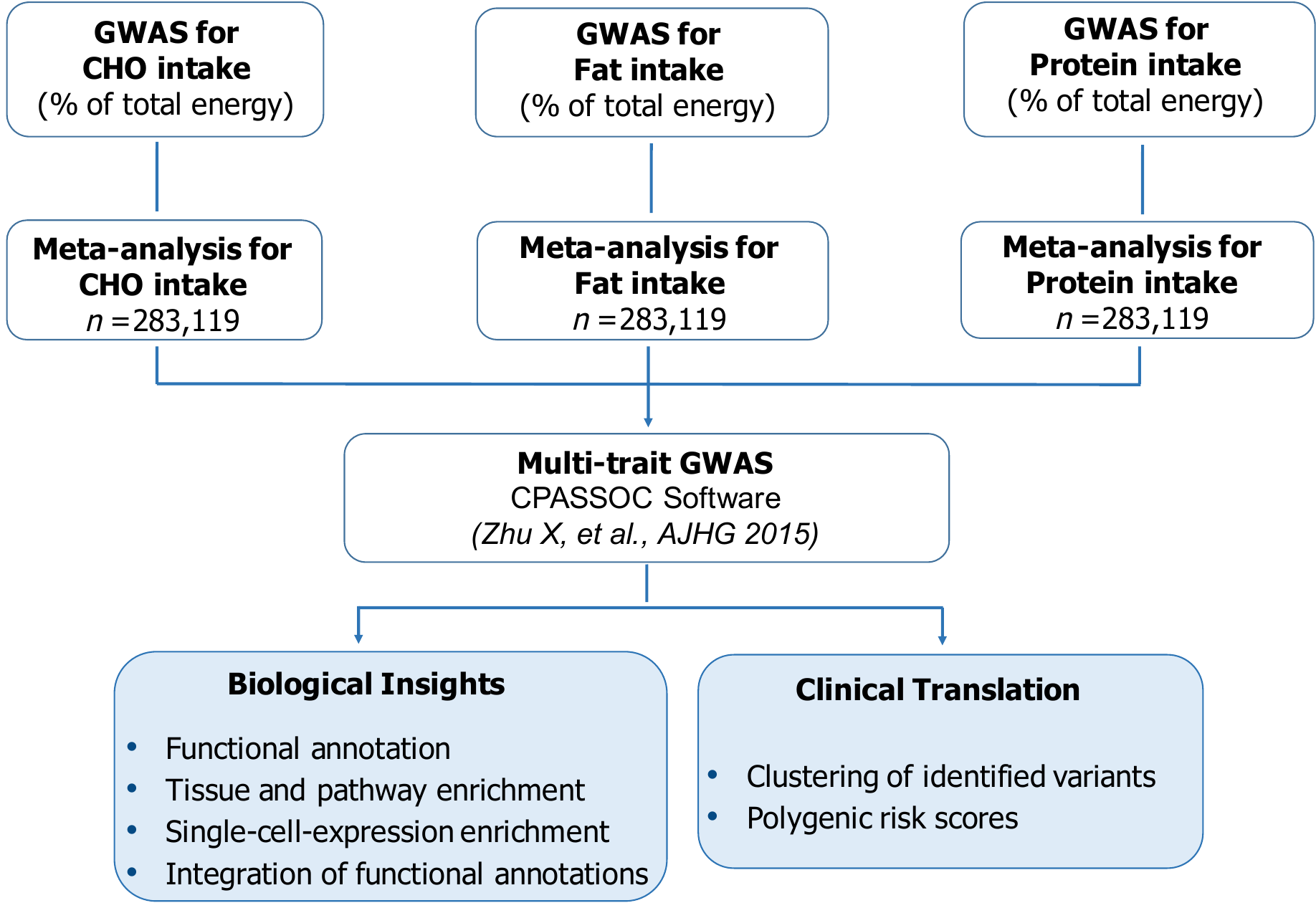

**Supplementary Fig. 1 | Schematic of the study design in the multi-trait genome-wide association meta-analysis for dietary intake in 283,119 individuals.** The genome-wide association meta-analysis of dietary intake comprised data from 192,005 participants from the UK Biobank and 91,114 participants from the CHARGE Consortium. Single-trait macronutrient GWAS from the UK Biobank and CHARGE Consortium were meta-analyzed using METAL and then combined into a multi-trait GWAS using the multi-trait CPASSOC. Downstream *in silico* analyses were conducted to identify biological features of identified loci including functional annotation, tissue and pathway enrichment, single-cell RNA expression analyses, and integration of functional annotations to re-weight the GWAS and identify new loci with high confidence of association. The Bayesian nonnegative matrix factorization clustering algorithm was used to classify dietary intake genetic loci into subgroups based on potential functional and clinical similarities. Cluster-based polygenic risk scores were built to investigate patterns of metabolic risk.

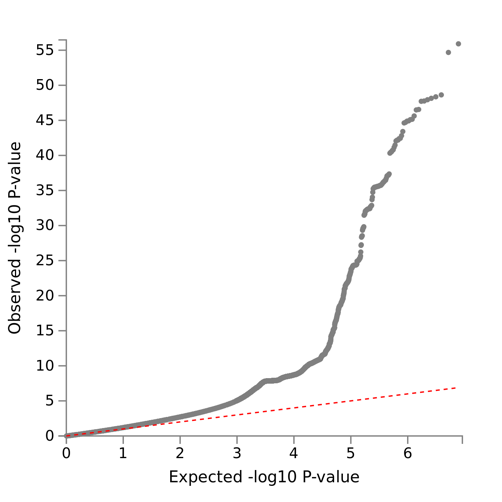

Multi-trait

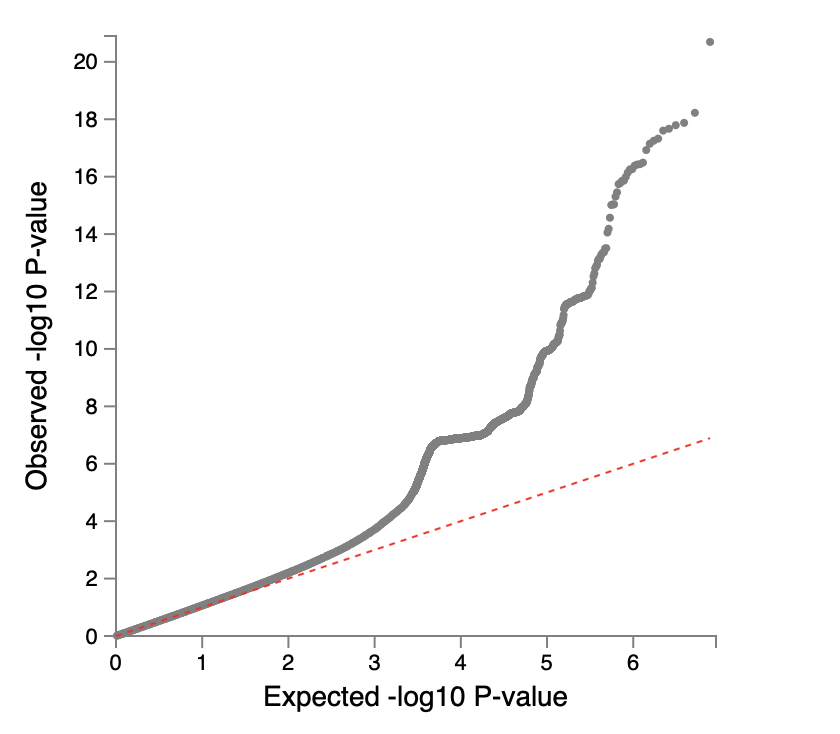

Carbohydrate intake

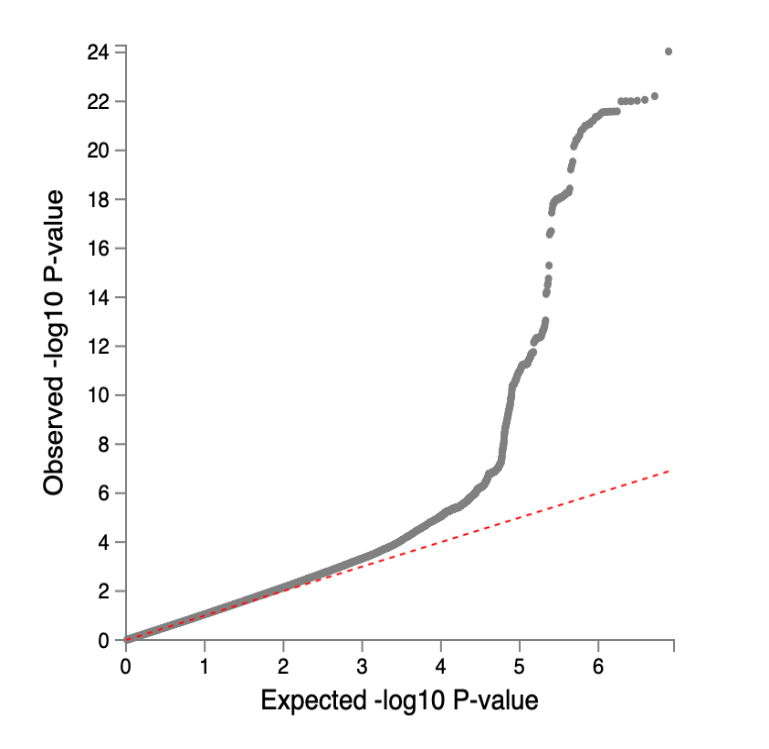

Fat intake

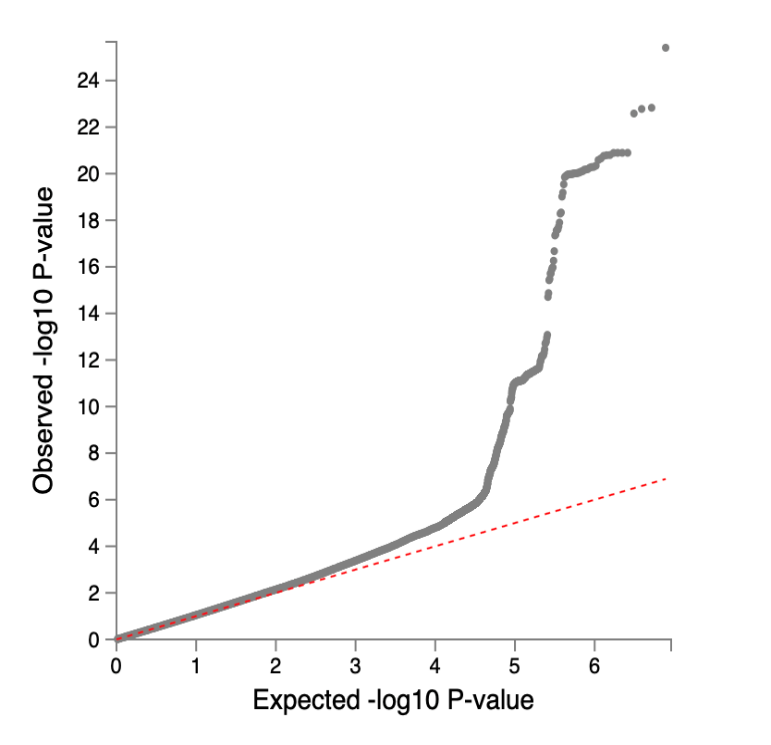

Protein intake

λ=1.13

Mean***𝜒*** ^2^=1.16

λ=1.14

Mean***𝜒*** ^2^=1.16

λ=1.17

Mean***𝜒*** ^2^=1.21

λ=1.15

Mean***𝜒*** ^2^=1.12

**d**

**c**

**b**

**a**

**Supplementary Fig. 2 | Quantile-quantile plot of the SNP-based associations with single-trait and multi-trait genome-wide association meta-analyses of 283,119 individuals.** Quantile–quantile plot of the SNP-based associations with multi-trait (a), carbohydrate (b), fat (c), and protein intake (d). SNP P values were computed in METAL by weighting effect size estimates using the inverse of the corresponding standard errors.

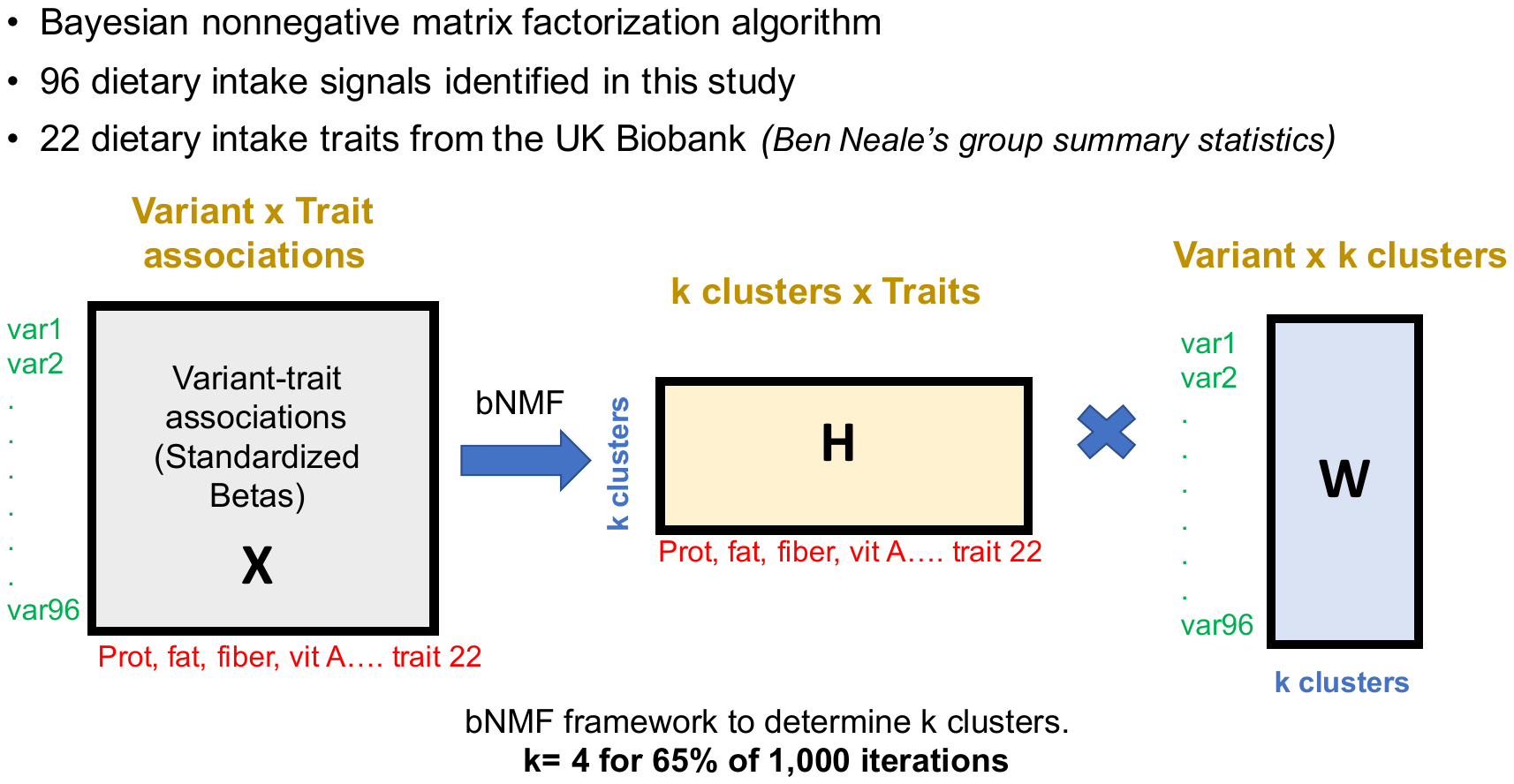

**k= 3 for 65% of 1,000 iterations**

79 genetic variants reaching nominal significance association with proportion fat intake

**Supplementary Fig. 3 | Schematic overview of the Bayesian nonnegative matrix factorization clustering algorithm.** The input for the Bayesian nonnegative matrix factorization clustering algorithm (bNMF) was the set of 79 genetic variants reaching nominal significance association with proportion fat intake. Next summary association statistics for 22 dietary intake traits from the UK Biobank were aggregated for each dietary intake variant. Our analyses involved variants aligned by their alleles associated with increased fat intake. We generated standardized effect sizes for variant trait associations from GWAS by dividing the estimated regression coefficient beta by the standard error, using the UK Biobank summary statistic results (variant-trait association matrix (79 by 22)). The defining features of each cluster were determined by the most highly associated traits, which is a natural output of the bNMF approach. bNMF algorithm was performed in R for 1,000 iterations with different initial conditions, and the maximum posterior solution at the most probable number of clusters was selected for downstream analysis.

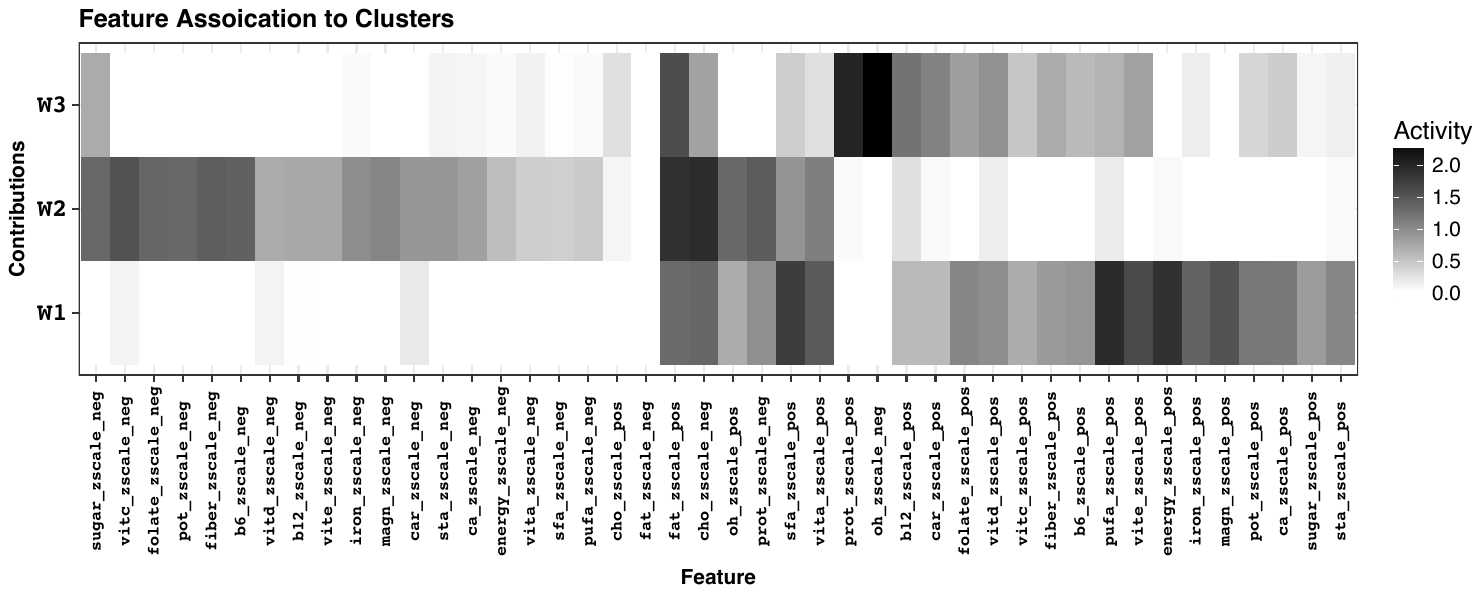

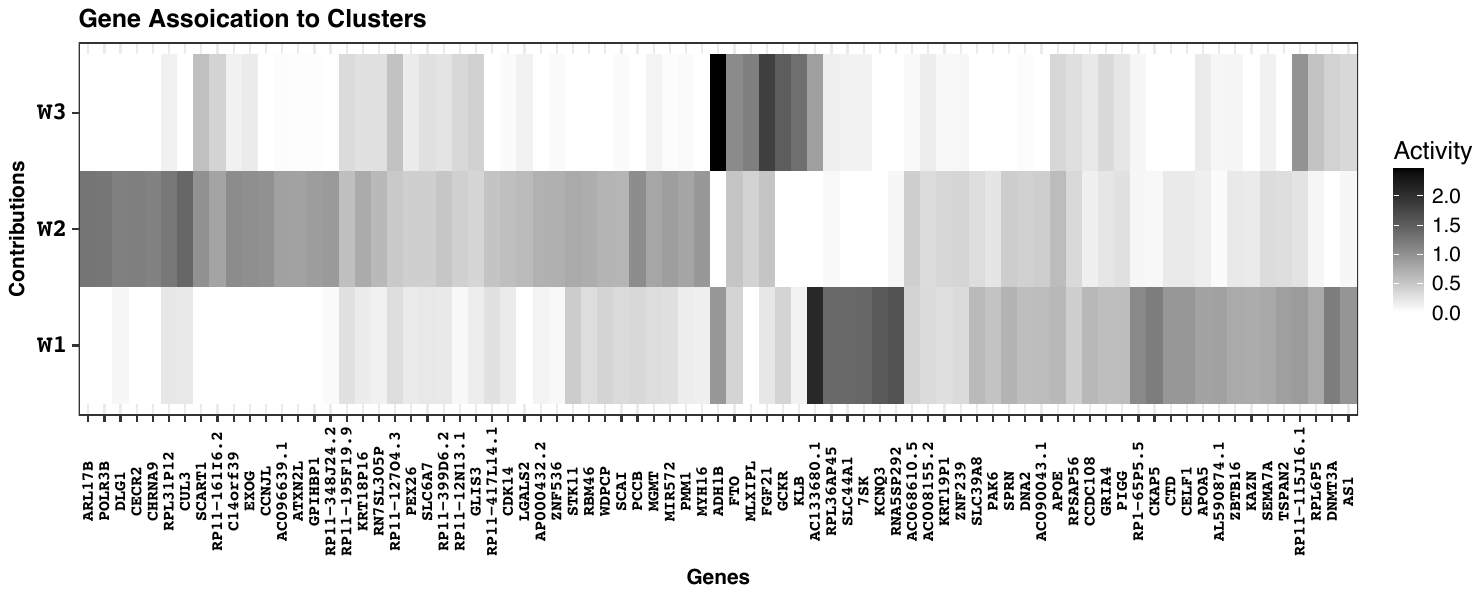

**Loci association to cluster**

**Trait association to cluster**

**a**

**b**

**Cluster-specific polygenic risk score and traits association**

**c**

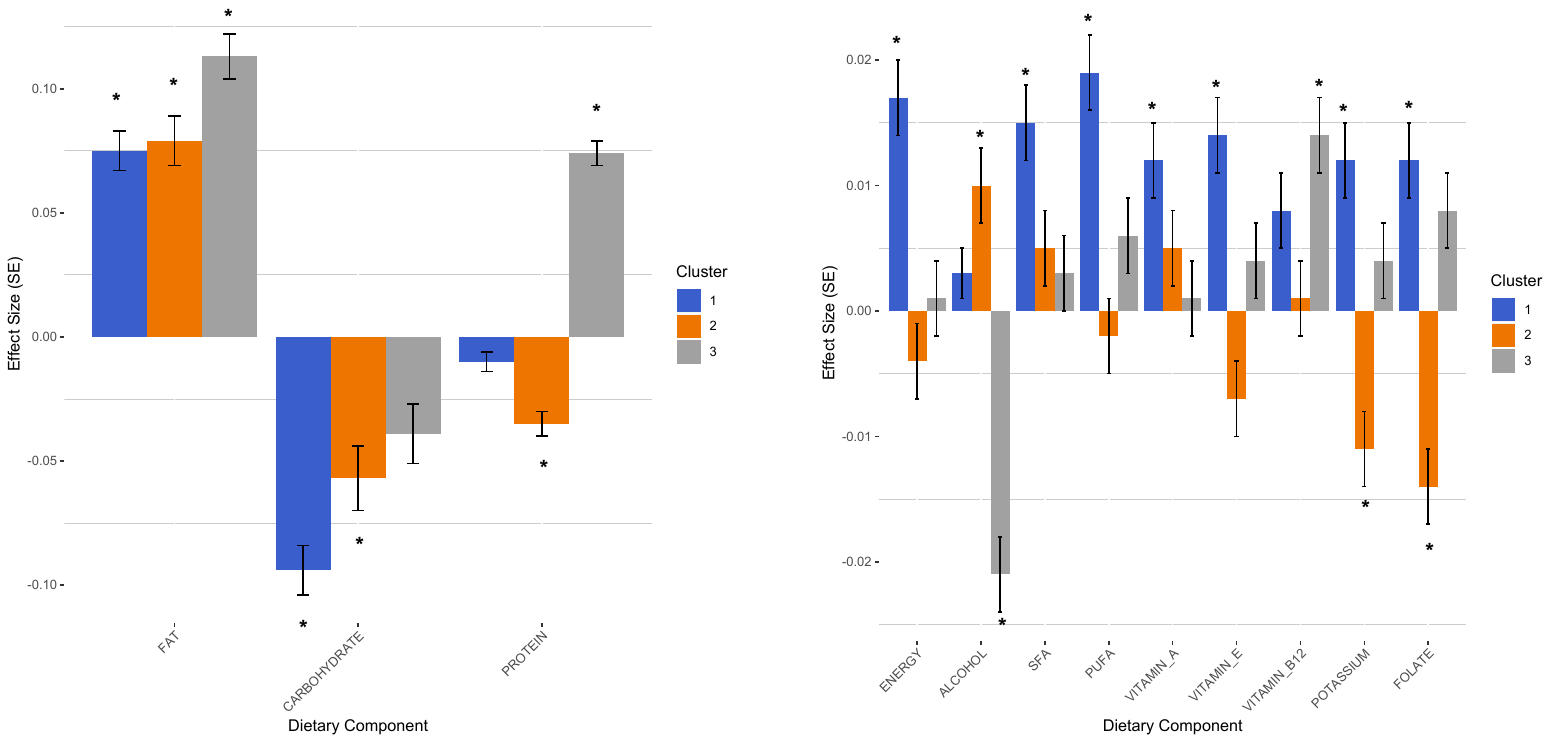

Weight

Weight

**Supplementary Fig. 4 | Trait and loci association to clusters.** Clustering of variant-trait associations was performed for 79 genetic variants reaching nominal significance association with proportion fat intake and 22 nutritional traits derived from GWAS using the Bayesian nonnegative matrix factorization clustering algorithm, with identification of four robust clusters present on 65% of iterations. Loci and traits defining each cluster were based on a cut-off of weighting of 1.09 (Methods). **a)** trait association to cluster, **b)** loci association to cluster:

*Cluster 1: AC133680.1, RNA5SP292, KCNQ3, X7SK, RPL36AP45, SLC44A1, CKAP5, DNMT3A*

*Cluster 2: CUL3, ARL17B, POLR3B, RPL31P12, CECR2, DLG1, CHRNA9*

*Cluster 3: ADH1B, FGF21, GCKR, KLB, MLXIPL*

**c)** To confirm that cluster-specific polygenic risk scores mirrored traits defining each cluster, we tested for associations of polygenic risk scores with each GWAS dietary trait using inverse-variance weighted fixed effects meta-analysis using summary statistics from the UK Biobank. Bonferroni threshold of significance for 22 traits and 3 clusters was set at 7.6×10^−4^ (displayed key traits).
